# Trophic status and local conditions affect microbial potential for denitrification versus internal nitrogen cycling in lake sediments

**DOI:** 10.1101/2021.07.12.452135

**Authors:** K.B.L. Baumann, R. Thoma, C.M. Callbeck, R. Niederdorfer, C.J. Schubert, B. Müller, M.A. Lever, H. Bürgmann

**Affiliations:** Eawag, Swiss Federal Institute for Aquatic Science and Technology, Department of Surface Waters-Research and Management, 6047 Kastanienbaum, Switzerland; ETH Zurich, Institute of Biogeochemistry and Pollutant Dynamics (IBP), Universitätstrasse 16, 8092 Zurich, Switzerland

**Keywords:** Metagenomics, microbial ecology, freshwater, porewater, DNRA, Nitrification, Denitrification, Anammox, Comammox

## Abstract

The nitrogen (N) cycle is of global importance as N is an essential element and a limiting nutrient in terrestrial and aquatic ecosystems. Excessive anthropogenic N fertilizer usage threatens sensitive downstream aquatic ecosystems. Although freshwater lake sediments remove N through various microbial transformation processes, few studies have investigated the microbial communities involved. In an integrated biogeochemical and microbiological study on a eutrophic and oligotrophic lake, we estimated N removal rates in the sediments from porewater concentration gradients. Simultaneously, the abundance of different microbial N transformation genes was investigated using metagenomics on a seasonal and spatial scale. We observed that contrasting nutrient concentrations in the sediments were reflected in distinct microbial community compositions and significant differences in the abundance of various N transformation genes. Within each lake, we observed a more pronounced spatial than seasonal variability. The eutrophic Lake Baldegg showed a higher denitrification potential with higher *nosZ* gene (N_2_O reductase) abundance and higher *nirS*:*nirK* (nitrite reductase) ratio, indicating a greater capacity for complete denitrification. Correspondingly, this lake had a higher N removal efficiency. The oligotrophic Lake Sarnen, in contrast, had a higher potential for DNRA and nitrification, and specifically a high abundance of *Nitrospirae*, including some capable of comammox. In general, the oligotrophic lake ecosystems had a higher microbial diversity, thus acting as an important habitat for oligotrophic microbes. Our results demonstrate that knowledge of the genomic N transformation potential is important for interpreting N process rates and understanding the limitations of the N cycle response to environmental drivers.

**Importance¶:** Anthropogenic nitrogen (N) inputs can lead to eutrophication in aquatic systems, specifically in N limited coastal ecosystems. Lakes act as N sinks by transforming reactive N to N_2_ through denitrification or anammox. The N cycle in lake sediments is mediated by microbial processes and affected by environmental drivers such as the amount and quality of settling organic material or nitrate concentration. However, the microbial communities mediating the different N transformation processes and their impact on N removal in freshwater lake sediments remain largely unknown. We provide the first seasonally and spatially resolved metagenomic analysis of the N cycle in the sediments of two lakes with different trophic states. We show that the trophic state of lakes provokes other microbial communities with characteristic key players and functional potential for N transformation.

## Introduction

Nitrogen (N) is an essential nutrient, and microbes play central roles in the natural N cycle. Only N-fixing microbes can convert di-nitrogen (N_2_) to reactive N (N_r_; N compounds readily available for biological conversion; e.g., NO_3_^-^, NO_2_^-^, NH_4_^+^, N_2_O). The different N_r_ compounds are used as electron acceptors or donors in several microbial metabolic pathways. Further, N can be a limiting nutrient in many terrestrial and aquatic ecosystems, such as the coastal ocean ^1–3^. Thus, excessive anthropogenic N_r_ input, e.g., from runoff of agricultural fertilizer, human wastewater, and fossil fuel combustion, can lead to trophic changes in aquatic ecosystems and the occurrence of harmful algal blooms ^4, 5^. Switzerland is an essential headwater system of European rivers and a non-negligible N source despite its small area. N loads from atmospheric deposition (44 kt yr^-1^), mineral fertilizers (52 kt yr^-1^), and sewage (43 kt yr^-1^) are about equally responsible for the N contamination of ecosystems in Switzerland ^6^. Atmospheric deposition rates on the Swiss Plateau, locally exceeding 40 kg N ha^-1^yr^-1^, are among the highest in the world ^6,7^. Today, Switzerland exports 63 kt yr^-1^ dissolved N via the Rivers Rhine and Rhone (^8^, an average of 1995-2013). This load corresponds to 1.3% of the total export of Europe to the seas (4761 kt yr^-1^; ^9^) and has remained high over recent decades.

Lakes have been identified as important N sinks that convert up to 90% of reactive N to less bioavailable N_2_ gas and thus substantially reduce N loads to oceans ^10^. N removal occurs through several microbially-mediated processes, mainly in the sediment at the oxic-anoxic transition zone. The main N loss processes are denitrification and anaerobic ammonia oxidation (anammox), both of which produce N_2_ as an end product, or long-term burial (sequestration) ^11–17^. The effectiveness of this removal process depends on interactions with other processes in the N cycle that provide or compete for their substrates and various other environmental factors. A better understanding of the environmental and ecological controls of these N removal processes is thus necessary to understand and predict the N removal by lakes today and under future conditions of global change.

Many different environmental drivers influence the N transformation processes in lakes. Suitable growth conditions and N substrates are key factors for the removal process. Accordingly, parameters that influence N removal rates are NO_3_^-^ concentration, organic matter (OM) input and remineralization, or redox conditions, indicated e.g., by O_2_ concentrations, among others ^1, 4, 10, 11, 16, 18–23^. Most of these factors are intricately linked to the overall trophic status of lakes. For example, Finlay et al. ^12^ reported seven-fold higher N removal rates in eutrophic than in oligotrophic lakes. Other studies found that inter- and intra-lake variations in denitrification rates were positively correlated with water residence times and N loads ^5, 24, 25^.

It is less well understood how these system observations are linked to the ecology of the microbial communities that mediate these processes. By studying the diversity, ecology, and identity of the microbial populations involved in the N transformation processes, we better understand the mechanisms underlying efficient N removal in lake ecosystems. Microbial ecology has made considerable progress over recent decades on detecting, characterizing, and quantifying the key gene families encoding for central enzymatic systems of the N transformation pathways (N transformation genes) in environmental systems using a variety of molecular tools ^15, 17, 26–29^. This has improved our understanding of the microbial ecology of nitrification, complete ammonia oxidation (comammox), assimilatory and dissimilatory nitrate reduction to ammonia (ANRA and DNRA), denitrification, anammox, and N_2_ fixation^1, 14, 22, 27, 28, 30, 31^. Metagenomic sequencing technology allows for a more holistic view on microbial N cycling by characterization of multiple N gene families in parallel ^32^. Several metagenomic studies focused on understanding the N cycle and the microbes involved. ^33^, for example, found a significant change in the microbial community and its metabolic function along a gradient of N availability in different soils. Nelsen et al. ^34^ showed that the soil C and N content explained the N pathway frequency and that N cycling specialists were encoding a few transformation processes, as well as generalists were encoding all N transformation processes. Furthermore, Graf et al. ^29^ reported that *nirS* (cytochrome cd_1_ variant for nitrite reductase) denitrifiers often contained *nor* and *nosZ* genes, encoding for the nitric oxide and nitrous oxide reductase, thus have the potential for the complete denitrification pathway comparing to the *nirK* (copper-binding variant of nitrite reductase) denitrifiers. ^35^ described atypical *nosZ* microbes from soil metagenomes. Additionally, metagenomes provided a comprehensive insight on N cycling in different natural ecosystems such as alpine or forest soil, grassland ecosystems, or the Arabian Sea oxygen minimum zone ^26, 36–38^. The recent collation of N gene families into a comprehensive database provided an improved framework for surveying N cycling processes in environmental metagenomes ^32^. Combined, more traditional molecular approaches and metagenomic investigations improved our understanding of the microbial N network in many stratified ecosystems with an oxic-anoxic transition zone such as open oceans or estuaries ^1–3, 39^. This indicates that this framework provides a promising approach also for freshwater sediment N cycling.

Our study investigated the microbial community structure in lake sediments of eutrophic Lake Baldegg and oligotrophic Lake Sarnen over seasonal and spatial scales. We aim to link characteristics of the microbial communities involved in N transformation processes with the porewater fluxes of nitrate and ammonium recently reported for the same lakes by Müller et al. ^20^. To investigate the microbial community composition over seasonal, locational, and sediment depth scales, we used a high throughput 16S rRNA gene amplicon sequencing approach. Furthermore, we sequenced shotgun metagenomes and applied advanced bioinformatics to provide a detailed view of N gene abundances and their phylogenetic identity. We hypothesize that the lakes’ contrasting N transformation processes are based on distinct microbial community composition and that the microbial N cycling potential is linked to environmental drivers such as different nutrient levels and OM inputs in the studied lakes. Furthermore, we suggest that the varying environmental conditions at different locations and depths of a lake result in significant seasonal and spatial differences of the microbial N transformation potential, thus questioning the common practice of relying on single core sampling to characterize whole lake ecosystems.

## Material and Methods

### Study sites

Lakes Baldegg and Sarnen are situated on the Swiss plateau and the peri-alpine region, respectively. Both are small lakes, with a surface of 7.15 km^2^ and a maximum depth of 49 m for Lake Sarnen and 5.22 km^2^ and 66 m for Lake Baldegg ^20^. Lake Sarnen is surrounded by mountains, forests, and extensive agriculture, and therefore remained oligotrophic throughout the 20th century, maintaining a total phosphorus (TP) concentration of ∼5 mg P m^-3^. Lake Baldegg, in comparison, receives high P loads due to pig and cattle farming in the catchment. In the second half of the 20^th^ century, this lake experienced pronounced eutrophication that peaked in the 1970s. To prevent bottom water anoxia, the lake is artificially aerated with O_2_ during the stratified season since 1984 ^40^. Today, the lake is still eutrophic with TP concentrations of ∼24 mg m^-3^ ^41, 42^. Sediment trap data from our parallel study in 2017 and 2018 found a higher sedimentation rate of N-enriched organic matter (OM) in Lake Baldegg (56.8 gC m^-2^ yr^-1^ and 11.2 gN m^-2^ yr^-1^, C:N 5.9) compared to Lake Sarnen (28.6 gC m^-2^ yr^-1^ and 4.1 gN m^-2^ yr^-1^, C:N 8.5) ^20^. Both lakes are dimictic with an oxic water column throughout the year; thus, the oxic-anoxic transition zone, where most of the N transformation processes occur, extends over the first few mm of the sediments. To study spatial variability, we selected five sampling locations representing different depths between the inflow and outflow of the lakes (Fig. 1). The seasonal dynamics were investigated by sampling each location in March, May, August, and September 2018 (for details, see Müller et al. ^20^). Sediment depth was resolved by analyzing porewater through sampling ports or cutting layers from the core, as described below.

**FIGURE 1:**
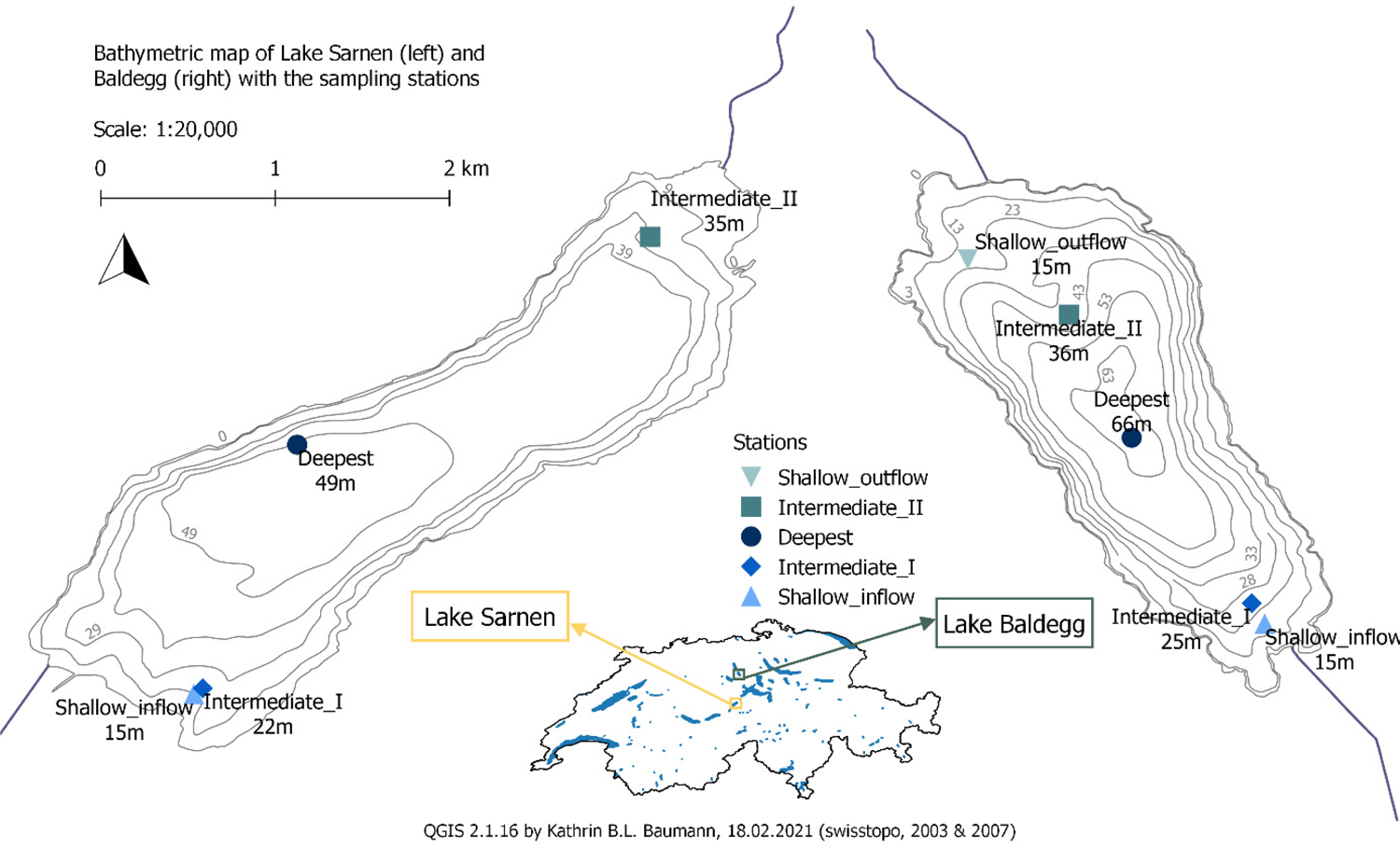
Sampling locations in Lake Sarnen (SAR) (left, yellow) and Lake Baldegg (BAL) (right, green), sampled in 2018 in March, May, August, and September. Porewater (0.25 cm resolution) and 16S rRNA gene sequencing data (0.5 cm resolution) were measured in all sampling points from 0-5 cm. DNA from the top 3 cm was pooled for Metagenome analysis of the Deepest location (SAR and BAL), Shallow_inflow (SAR), and Shallow_outflow (BAL) in all months. Shallow_inflow of Lake Baldegg was additionally sampled in August and September. Sediment traps were deployed at the deepest point of each lake (dark blue), see Müller et al. ^20^.

### Sediment sampling, chemical, and statistical analysis

The sediment sampling, porewater analyses, and oxygen measurements are described in Müller et al. ^20^. Briefly, we used a gravity corer for the sediment sampling with PVC tubes (5.9 cm diameter and 60 cm length, Uwitec, Austria). The PVC tubes had pre-drilled ports to extract porewater samples (∼200 µl) in 0.25 cm depth resolution using MicroRhizon filter tubes (Rhizosphere Research Products, Wageningen, Netherlands; 0.2 µm pore size; 0.8 mm diameter). The porewater was sampled onshore immediately after core retrieval, transferred to 1.5 ml tubes, stored on ice in the dark, and analyzed within 24 h. NO_3_^-^, NO_2_^-^, SO_4_^2-^ and NH_4_^+^ were analyzed using two ion chromatography devices (cations: 882 Compact IC plus, anions: 881 Compact IC pro, Metrohm, Switzerland). In addition, vertical O_2_ concentration profiles were recorded with an O_2_ micro-optode mounted to an automated micromanipulator (Presens, Germany) immediately after core retrieval. The O_2_ and porewater profiles were visualized using ggplot2, and the spatial and temporal differences were calculated using ANOVA from the base statistical packages in R (Version 3.6.1) ^43, 44^.

### DNA sampling and extraction

At each sampling event, a sediment core was retrieved for DNA analysis. The core was cut with a metal slicing device into the following sections: 0-0.5 cm, 0.5-1 cm, 1-1.5 cm, 1.5-2 cm, 2-3 cm, 3-5 cm. The slicing device was washed with 80% EtOH and molecular grade water (DNAse/RNAsefree, SigmaAldrich) before each core sampling and with molecular grade water during the slicing of cores. Three subsamples (∼2 ml) were taken from each section with a 3 ml syringe and immediately fixed in 2 ml RNAlater (Sigma Life Science) in a 5 ml tube (Qiagen, DNAse/RNAsefree). The sediment samples were kept at 4 °C until the next day and then stored at −80 °C. Prior to nucleic acid extraction, samples were thawed and washed three times with 2.5 ml 1x TE Buffer to remove excess salt from RNA Later. Nucleic acids were then extracted using the RNeasy PowerSoil total RNA kit in combination with the RNAeasy PowerSoil DNA Elution kit (Qiagen) according to the manufacturer’s instructions. Due to low RNA yields from Lake Sarnen, we only used the DNA for the analysis presented in this paper. The DNA quality and quantity were measured with a Nanodrop spectrophotometer (Nanodrop^TM^ One), and extracts were stored at −80°C until sequencing. Extraction blanks were prepared to check for contamination during the extraction and used as negative controls in sequencing.

### Amplicon sequencing, pipeline, and statistical analysis

All amplicon sequencing was performed by Novogene (Hong Kong) following standard sequencing and quality control protocols. Extracted DNA from all stations and all seasons (120 samples from Lake Baldegg and 96 samples from Lake Sarnen) was amplified with universal 16S rRNA gene primers 341F (5’-CCT AYG GGR BGC ASC AG-3’) and 806R (5’-GGA CTA CNN GGG TAT CTA AT-3’) targeting the V3-V4 region (Fragment length ∼466 bp), and sequenced with Illumina HiSeq technology to generate paired-end reads. This resulted in 119 samples with an average of 112,857 raw paired-end reads in Lake Baldegg, and 90 samples with an average of 139,822 raw paired-end reads in Lake Sarnen. The sequences were analyzed separately for each lake using the DADA2 pipeline in R (version 3.5.1) ^44, 45^. Briefly, the raw sequence adapters were trimmed, dereplicated and error rates were calculated before applying the DADA2 core sample inference algorithm. The forward and reverse reads were merged to obtain the amplicon sequence variants (ASVs). Following chimera removal with the removeBimeraDenovo algorithm, the taxonomic assignment of the ASVs was performed using a Naïve Bayesian classifier method based on the SILVA database 132 for V3/V4 region ^46^. Unclassified and rare ASVs (less than five instances in at least 10% of all samples) were filtered out. The ASV abundance tables and the corresponding environmental parameters were further analyzed by calculating the alpha and beta diversity (RDA=redundancy analysis), testing for significantly different microbial community composition between the lakes, locations, sampling months, and sediment depth using adonis and visualized with the phyloseq, ggplot2, vegan and microbiomeSeq packages using R statistical software (Version 3.6.3) ^43, 44, 47–49^. Details of the 16S rRNA gene amplicon library and pipeline can be found in the supplements (Table S1) and on GitHub (page).

### Metagenomic sequencing, pipeline, and statistical analysis

For metagenomic sequencing, DNA extracts from all samples of one core were pooled, using 1 µg of DNA from each sampling depth (0 –0.5 cm, 0.5-1 cm, 1-1.5 cm, 1.5-2 cm, and 2-3 cm) after measuring the DNA concentration with Qubit (QubitⓇ 2.0 Fluorometer, Invitrogen). We obtained eight metagenomes from Lake Sarnen (deepest and shallow_inflow station for March, May, August, and September) and ten metagenomes from Lake Baldegg (deepest and shallow_outflow locations for March, May, August, and September, and shallow_inflow for August and September). The metagenomes were sequenced using the Illumina NextSeq platform to generate 150 bp paired-end reads (averaging ∼350 bp in length) by Novogene (Hong Kong). The quality of each metagenome was evaluated using FastQC ^50^. Prinseq was used to trim the metagenomics reads (minimum quality mean 20) ^51^. Each sample was assembled using Megahit with the options meta-sensitive and –min_contig length 500 ^52^ and reads were mapped back using BBmap ^53^. The sam files were converted to bam files using sambamba ^54^ and SAMtools ^55^ and the reads were subsequently counted using featureCounts from subread 1.6.4 ^56^. For functional annotation, we used prokka v1.13 ^57^. The N transformation genes (Table 1) were obtained by mapping the protein output from prokka against the N cycle database (NCycDB; accessed July 2019), applying a 100% identity criterion ^32^. The genes and representative gene families were assigned to N cycle processes based on the information provided by NCycDB (Table 1 in Tue et al. (2018)); if the gene was assigned to more than one process in this table, we made a new cluster named by the processes (e.g., Denitrification_DNRA) (Table 1). All further analysis and visualization were done in R (Version 3.6.3) ^44^. The quantified genes were normalized for the gene lengths and sequencing depth by calculating genes per million (gpm) for the visualization. We calculated the *nirS:nirK* ratio from the gpm data to indicate the ratio of complete denitrifiers vs. incomplete denitrifiers (Graf et al., 2014). The principal component analysis and testing for significantly different N transformation genes between lakes (alpha=0.001) were calculated using DESeq2 ^58^. We used the non-normalized count files from feature counts as DESeq2 has a normalization step included (following the workflow from ^59^). We used the wald test, with the condition Lake Baldegg vs. Lake Sarnen, to extract the significantly different N transformation genes using the Benjamini & Hochberg method from DESeq2 to correct p-values for multiple testing ^58, 59^. The results were visualized with ggplot2 and ComplexHeatmap ^60^ after calculating the z-score (z=(x-mean(x))/sd(x), whereas x=N gene counts; sd=standard deviation) for all significantly different N transformation genes to emphasize the differences. Finally, candidate key players for the significantly different genes and *nirS* were resolved by taxonomic assignment of the prokka output using Kaiju ^61^. Kaiju is a taxonomic assignment tool developed for short reads and may be biased for long reads such as assembled contigs. Kaiju still contains the former class Betaproteobacteria, which we assigned to Gammaproteobacteria as in SILVA database 132.

**TABLE 1:**
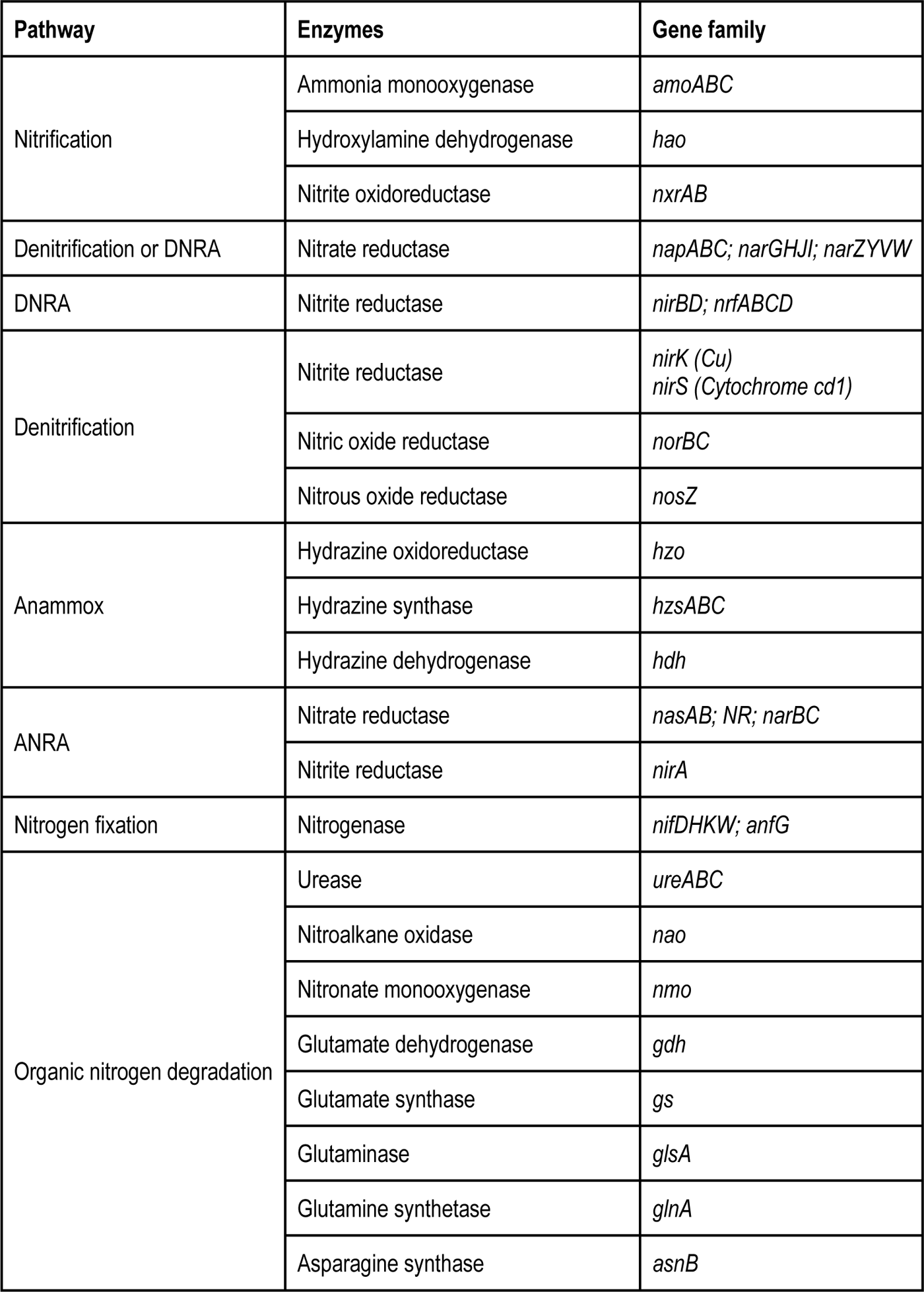
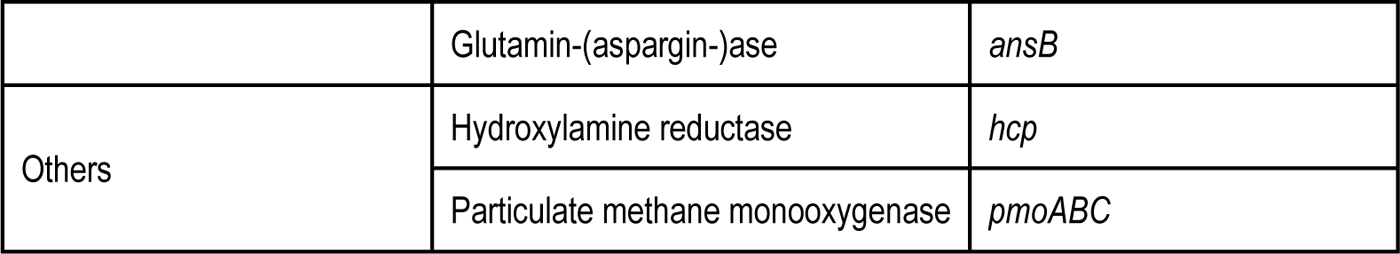
Overview of the nitrogen transformation pathways, processes, and the corresponding gene families, which were analyzed using NCycDB (adapted from Tu et al. ^32^). Abbreviations stand for ANRA: assimilatory nitrate/nitrite to ammonium; DNRA: dissimilatory nitrate reduction to ammonium; Anammox: anaerobic ammonium oxidation.

We tested the spatial and temporal N transformation differences with DESeq2 using lakes + Station and lakes + months as conditions. The results revealed 14 significantly different genes between the lakes and sampling location (p<0.01) and 0 significantly different genes between the lakes and sampling time point (p<0.01).

The sequencing depth, quality, filtered reads, mean contig length, and mapped reads for each sample are summarized in the supplementary table (Table S2).

### Data availability

All sequences can be found under the NCBI BioProject PRJNA726540. The 119 16S rRNA gene sequences and the metadata from Lake Baldegg have the accession numbers SAMN18962412-SAMN18962530. The 90 16S rRNA gene sequences and the metadata of Lake Sarnen are deposited under the accession numbers SAMN18978354-SAMN18978443. The ten metagenomes and the metadata can be found under the accession numbers SAMN19067306-SAMN19067315. The eight metagenomes and the metadata from Lake Sarnen can be found under the accession numbers SAMN19030157-SAMN1903164. All other data will be published after acceptance of the manuscript at the institutional data repository of Eawag (ERIC) (https://data.eawag.ch/), following FAIR data sharing principles.

## Results

### Microbial community composition and environmental drivers

Sediment porewater profiles from both lakes showed pronounced gradients of electron acceptors and reduced compounds, typical of freshwater lake sediment underlying an oxygenated water column. The sediment oxygen concentrations at the sediment-water interface were ∼20 times higher in Lake Sarnen (max 217 µMol) compared to Lake Baldegg (max 25 µMol) (Fig. 2A and F). O_2_ also penetrated up to 4 times deeper in Lake Sarnen (< 2 cm) than in Lake Baldegg (< 0.75 mm) (Fig. 2A and F). In both lakes, the highest O_2_ concentrations were measured at the shallower locations. O_2_ concentrations decreased in both lakes over the course of the year, with an average of 63 µMol in spring and 0 µMol in late summer in Lake Baldegg and 181 µMol in March and 67 µMol in September in Lake Sarnen. The NO_3_^-^ levels in Lake Baldegg were slightly higher (max 64 µMol) compared to Lake Sarnen (max 51 µMol) (Fig. 2B and G). In Lake Sarnen NO_3_^-^ penetrates deeper (< 4 cm), whereas NO_3_^-^ is depleted within the first 2 cm in Lake Baldegg. We did not observe any seasonal nor locational patterns for NO_3_^-^ (Fig. 2B and G). NO_2_^-^ concentrations generally remained below 2 µMol in both lakes (Fig. 2C and H). In Lake Baldegg, NO_2_^-^ was highest in the surface sediment (mean 0.21 µMol), in contrast to Lake Sarnen, where NO_2_^-^ peaked at 0.75 cm depth (mean 0.13 µMol) Fig. 2C and H). NH_4_^+^ concentrations increased with sediment depth in both lakes, but in Lake Baldegg (average 337 µMol) concentrations were over ten times higher compared to Lake Sarnen (average 21 µMol) (Fig. 2D and I). The NH_4_^+^ concentration in Lake Baldegg decreased from inflow to outflow (Fig. 2D). In Lake Sarnen, the deepest station showed the highest NH_4_^+^ concentration in the upper sediment (Fig. 2I). Lake Sarnen had a higher concentration of SO_4_^2^ than Lake Baldegg, with a mean SO_4_^2-^ concentration of 295 µMol and 27.5 µMol respectively (Fig. 2E and J). The SO_4_^2-^ depletion curve in Lake Baldegg was steeper and stabilized at a low concentration below 2 cm sediment depth, in contrast to Lake Sarnen where high SO_4_^2-^ concentrations were still measured at 4 cm sediment depth (Fig. 2E and J).

**FIGURE 2:**
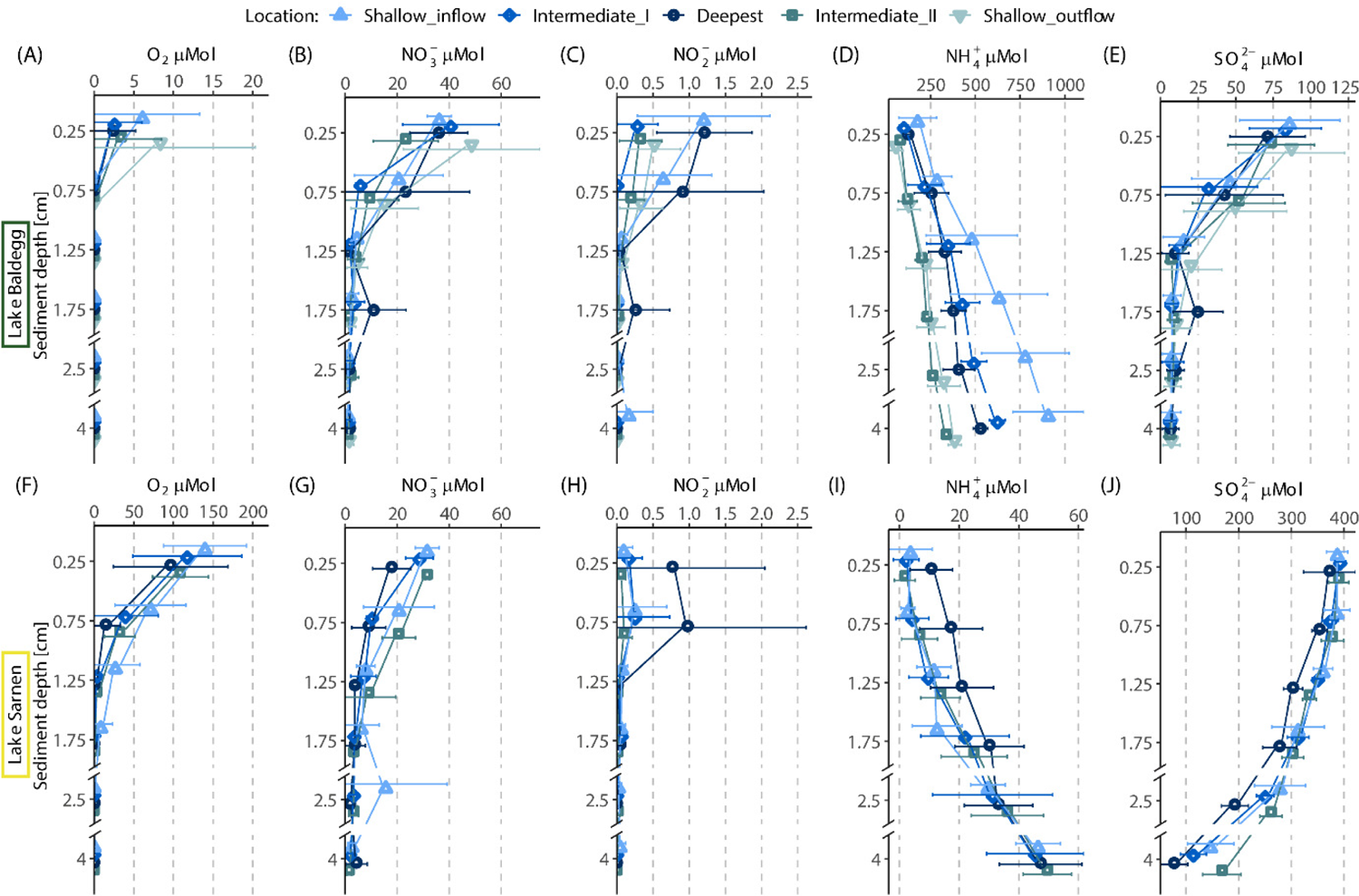
Porewater and oxygen profiles of Lake Baldegg (A-E) from the five different sampling locations (Shallow_inflow 15 m, Intermediate_I 25 m, deepest 66 m, Intermediate_II 36 m, and Shallow_outflow 15m) and Lake Sarnen (F-J) four different sampling locations (Shallow_inflow 15 m, Intermediate_I 22 m, deepest 49 m, and Intermediate_II 35 m). Each graph shows the seasonal means (points; the points from one sediment depth were plotted shifted to prevent overlay and to distinguish them better) with standard deviation (vertical error bars) for each location within sediment depth for the following chemical parameters: O_2_ (A and F), NO_3_^-^ (B and G), NO_2_^-^ (C and H), NH_4_^+^ (D and I) and SO_4_^2-^ (E and J).

We observed distinct microbial community composition and differences in microbial diversity in the two lakes. In Lake Sarnen, alpha diversity was significantly higher (Wilcoxon-test, p < 0.001) with an observed overall richness of 64,699 ASVs (unfiltered) compared to Lake Baldegg with 29,457 ASVs (Fig. S1A and B). The difference was robust to filtering rare ASVs (taxa appearing less than five times in at least 10% of the samples), leaving 2,599 ASVs for Lake Baldegg and 6,124 ASVs for Lake Sarnen (Fig. S1A and B).

The most abundant ASVs (top 500 ASVs) in Lake Baldegg were dominated by representatives of the phylum *Bacteroidetes* with *Bacteroidia* (31 %) and *Ignavibacteria* (6 %), *Gamma*-(26 %) and *Deltaproteobacteria* (16 %), *Euryarchaeota* with *Methanomicrobia* (5 %) and *Firmicutes* with *Clostridia* (2 %) (Fig. 3A). In Lake Sarnen, the phylum *Proteobacteria* was also abundant, but with a greater contribution of *Gamma*-(41 %) and *Alphaproteobacteria* (11 %). The *Bacteroidetes,* primarily of the class *Bacteroidia* (9 %), were three times less abundant than in Lake Baldegg. The third most abundant phylum in Lake Sarnen was *Nitrospirae* with genus *Nitrospira* (8 %), *Verrucomicrobiae* (8 %), and *Acidobacteria* (5 %) (Fig. 3B) which were only a minor component in Lake Baldegg. In Lake Sarnen, *Thaumarchaeota* (< 1 %) were the only archaeal phylum represented.

**FIGURE 3:**
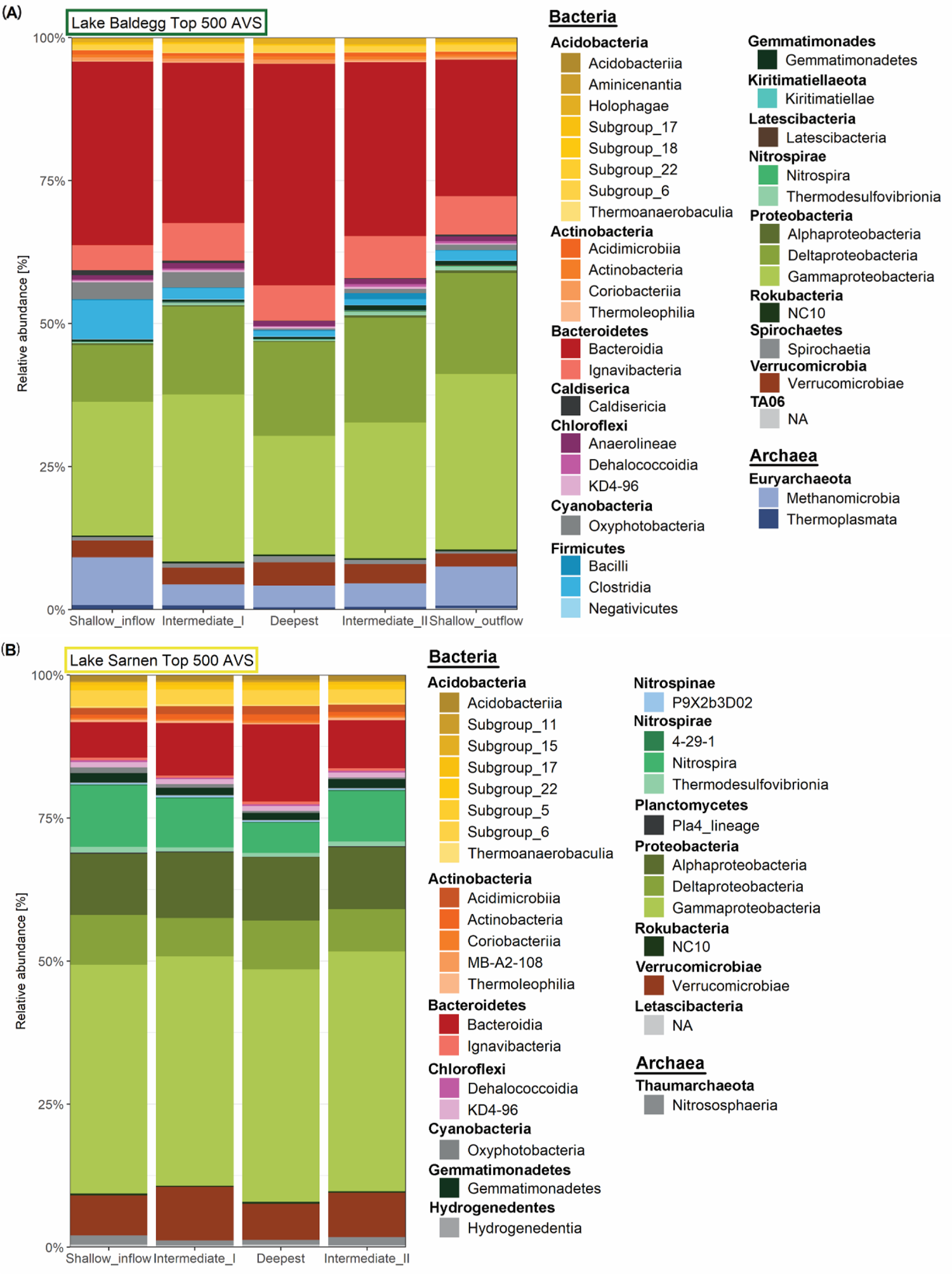
Microbial community composition of the top 500 ASVs on class level for Lake Baldegg (top) and Lake Sarnen (bottom). Classes that belong to the same Phylum are grouped together, and the Phyla is written above each class group. The data shown are averaged over sediment depth and time for each station per lake separately. The ASVs were retrieved with DADA2 and assigned using the SILVA132 database.

Constrained ordination by RDA confirmed that community composition was significantly different in the two lakes (PERMANOVA, p<0.001, R^2^=0.59) (Fig. S1C). The analysis also showed that beta-diversity patterns in the two lakes differed considerably (Fig. 4). In Lake Baldegg, the first ordination axis (CAP, 48%) reflects the location, and the second component (CAP2 25%) explains the microbial community composition differences with sediment depth (Fig. 4A, Fig. S2A and C). The microbial community composition showed a significant difference by sampling location (PERMANOVA, p<0.001, R^2^=0.44) in Lake Baldegg, changing from inflow to outflow (Fig. 4A). The influence of sediment depth and sampling month on the community was also significant (PERMANOVA, p<0.001), although with much lower explanatory power (R^2^=0.06 months and R^2^=0.05 sediment depth). In contrast, for Lake Sarnen, the first component reflected the significant microbial community composition variance with sediment depth (CAP1, 56%), and the second component the community differences with location (CAP2, 18%), indicating fewer differences between locations than in Lake Baldegg (Fig. 4B). The community composition varied significantly between locations in Lake Sarnen (PERMANOVA, p<0.001, R^2^=0.21). The deepest location of Lake Sarnen revealed the most distinct community composition compared to the three other locations, where the two Intermediate stations showed the most similar microbial community composition (Fig. 4B). As in Lake Baldegg, the community differences by sediment depth and sampling month were significant but with lower explanatory power (R^2^=0.09 sediment depth and R^2^=0.05 months) (Fig. S2B and D). RDA indicated that NH_4_^+^ concentration displayed the highest correlation with the observed changes in community composition in both lakes’ deeper sediments (Fig. S2A and B). In contrast, the other four environmental drivers (O_2_, NO_3_^-^, NO_2_^-^ and SO_4_^2-^) were drivers for the microbial communities living in the surface sediment (Fig. S2A and B).

**FIGURE 4:**
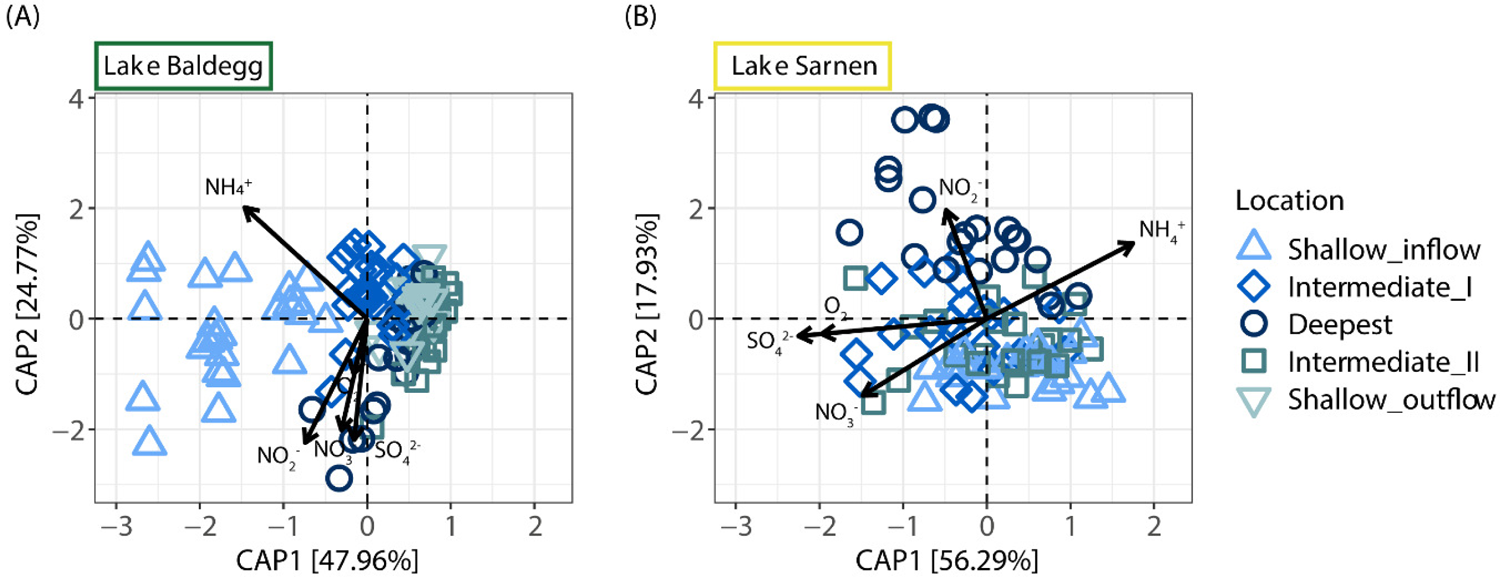
Constrained ordination biplots (Redundancy analysis (RDA) on weighted UniFrac distance) of microbial community composition and the different environmental drivers (O_2_, NO_3_^-^, NO_2_^-^, NH_4_^+^ and SO_4_^2-^) for Lake Baldegg (A) and Lake Sarnen (B). In Lake Baldegg the two main components (CPA1 and CPA2) explain 72% of the differences in the microbial communities. In Lake Sarnen the main components (CPA1 and CPA2) explain 74 % of the differences in the microbial communities. The samples are colored and shaped by location.

### Microbial nitrogen transformation potential and possible key players

A total number of 14,976 and 12,712 genes were assigned to genes indicative of N transformation processes in Lake Baldegg and Sarnen, respectively (Table S2). The N transformation genes in Lake Baldegg were assigned to N degradation and biosynthesis (34,355 gpm), DNRA (7,168 gpm), ANRA (4,886 gpm), nitrate reduction genes belonging to either DNRA or denitrification (4,069 gpm) or specific for denitrification (4,134 gpm), N_2_ fixation (1,118 gpm), nitrification (358 gpm), and anammox (104 gpm) (Fig. 5A, Table S3). Hydroxylamine reductase (*hcp*) was relatively abundant (1,000 gpm). In Lake Sarnen N degradation (26,612 gpm) and specific denitrification genes (4,269 gpm) were likewise abundant. But, we observed markedly fewer DNRA (4,738 gpm), N_2_ fixation (432 gpm), anammox (7 gpm), and hcp genes (86 gpm), and, in contrast, a greater abundance of nitrification (1,162 gpm), denitrification, and nitrate reduction genes (4,306 gpm), than in Lake Baldegg (Fig. 5B, Table S3). As a non-nitrogen cycle reference gene, we further analyzed methane-monooxygenase (*pmoABC*), which were in both lakes less abundant (Lake Baldegg 116 gpm and Lake Sarnen 283 gpm) (Table S3). N transformation gene composition showed the most vital trends in relation to lakes (PC1, 62%) and locations (PC2, 8%) (Fig. 5C), whereas seasonal variations were comparatively minor (Fig. 5C, M&M section).

**FIGURE 5:**
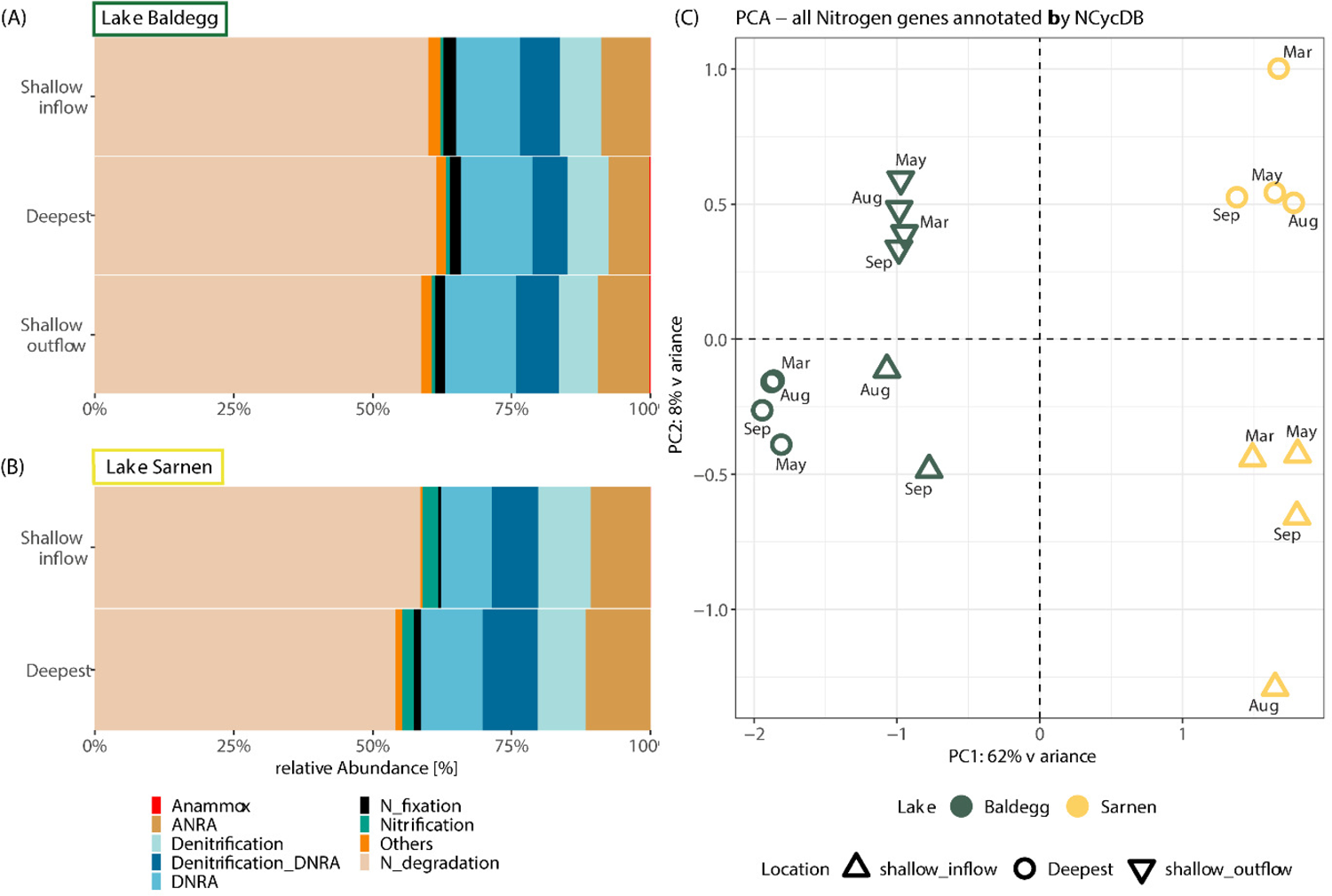
Nitrogen transformation gene distribution in Lake Baldegg (A) and Lake Sarnen (B) for each station summed over all seasons, grouped by the nitrogen transformation processes (Tab. 1). Nitrate reduction genes that can not be assigned to either Denitrification or DNRA are grouped together (Denitrification_DNRA). The principal component analysis (C) was done with the nitrogen gene counts for each station, calculated with the DESeq2 package.

We found 14 genes involved in various N transformation pathways that had significantly different abundances between Lake Baldegg and Lake Sarnen (Deseq2, Wald test, padj<0.001) (Fig. 6). As for the microbial community composition, we did not observe a distinct temporal clustering but a clear spatial clustering in the heatmap (Fig. 6). Lake Sarnen had a significantly higher abundance of bacterial ammonia (*amoABC_B*) and nitrite-oxidizing (*nxrB*) genes than Lake Baldegg (Deseq2, Wald test, padj < 0.001), indicating a higher nitrification potential (Fig. 6). In Lake Baldegg, the shallow outflow location showed a higher nitrification potential than the deep location (Fig. 6). Lake Baldegg had a significantly higher potential to reduce NO_3_^-^ via periplasmic nitrate reductase (*napA*) (Deseq2, Wald test, padj<0.001), whereas Lake Sarnen showed a significantly higher abundance for a different nitrate reductase gene, namely *narH* (Deseq2, Wald test, padj<0.001) (Fig. 6). Two representative genes of the DNRA step (*nrfC, nrfD*) encoding for NO_2_^-^ reduction to NH_4_^+^ were significantly higher in Lake Baldegg than in Lake Sarnen (Deseq2, Wald test, padj<0.001), for (Fig. 6). Among denitrification genes, *nirK* was significantly more abundant in Lake Sarnen, especially in the shallow location (Deseq2, Wald test, padj<0.001), while *nosZ* was significantly more abundant in Lake Baldegg (Deseq2, Wald test, padj<0.001). The other nitrate reductase gene, *nirS,* was similar in both lakes, resulting in a higher *nirS*:*nirK* ratio for Lake Baldegg than Lake Sarnen, especially in the shallower locations (Fig. 7). Anammox genes were significantly more abundant in Lake Baldegg (Deseq2, Wald test, padj < 0.001), and were highest in the shallow outflow location(Fig. 6). The assimilatory NO_3_^-^ reduction to NO_2_^-^ potential (*nirA*) was significantly higher in Lake Sarnen compared to Lake Baldegg (Deseq2, Wald test, padj < 0.001) (Fig. 6). In contrast, Lake Baldegg showed a significantly higher abundance of *nifHK*, which encodes for nitrogenase, the central enzyme for N_2_ fixation. The N_2_ fixation potential was especially high in the deep and shallow inflow locations (Deseq2, Wald test, padj < 0.001) (Fig. 6). The hydroxylamine reductase gene *hcp* was significantly more abundant (Deseq2, Wald test, padj < 0.001) in Lake Baldegg.

**FIGURE 6:**
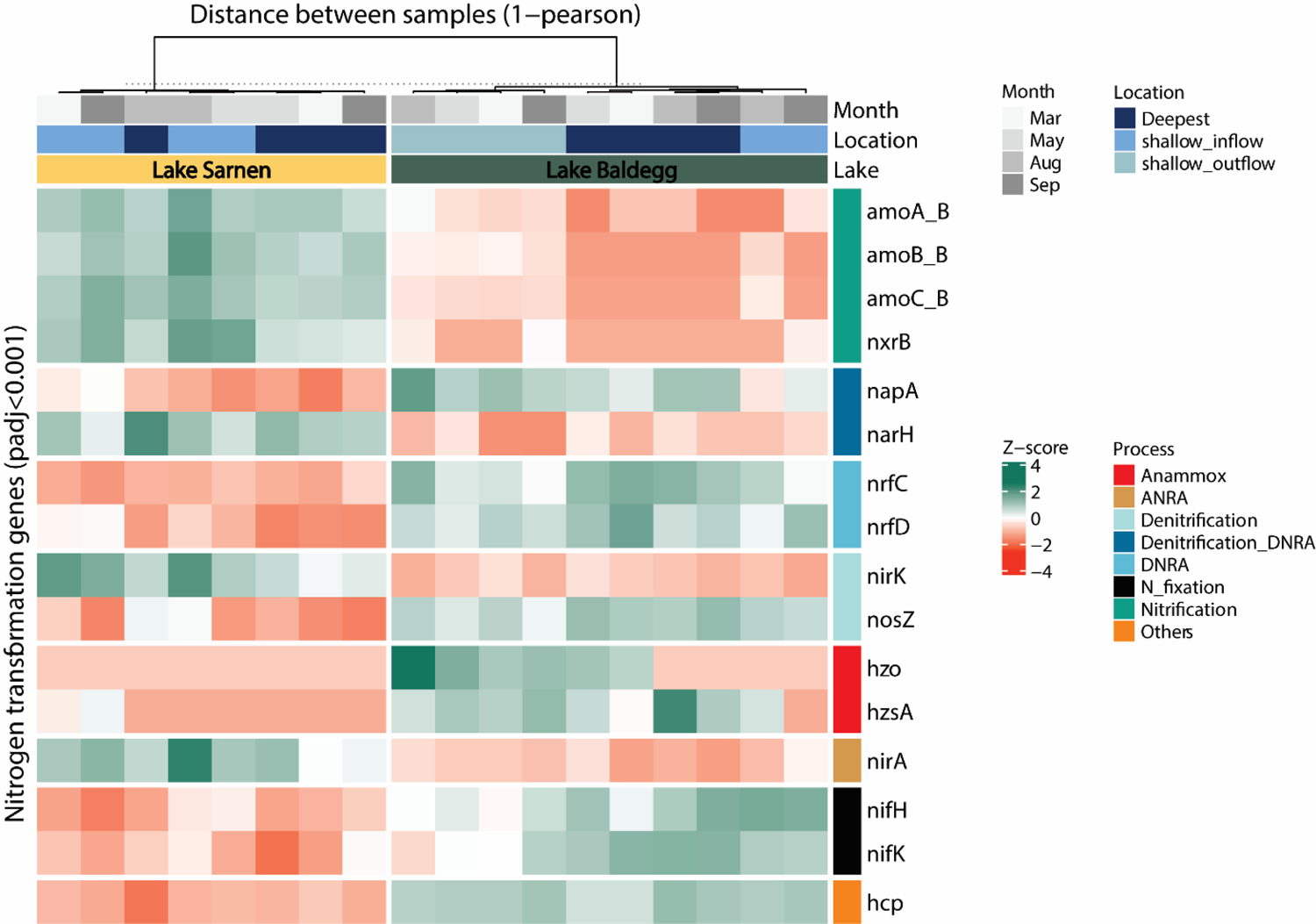
Heatmap of nitrogen gene abundances with significant differences between Lake Baldegg and Sarnen (alpha=0.001, padj=adjusted p-value with Benjamini&Hochberg method from DESeq2) clustered by Lake and station (column clustering distance calculated using Pearson in ComplexHeatmap) in the columns and nitrogen process in the rows. The z-score of abundances was calculated for each gene individually to emphasize the differences.

**FIGURE 7:**
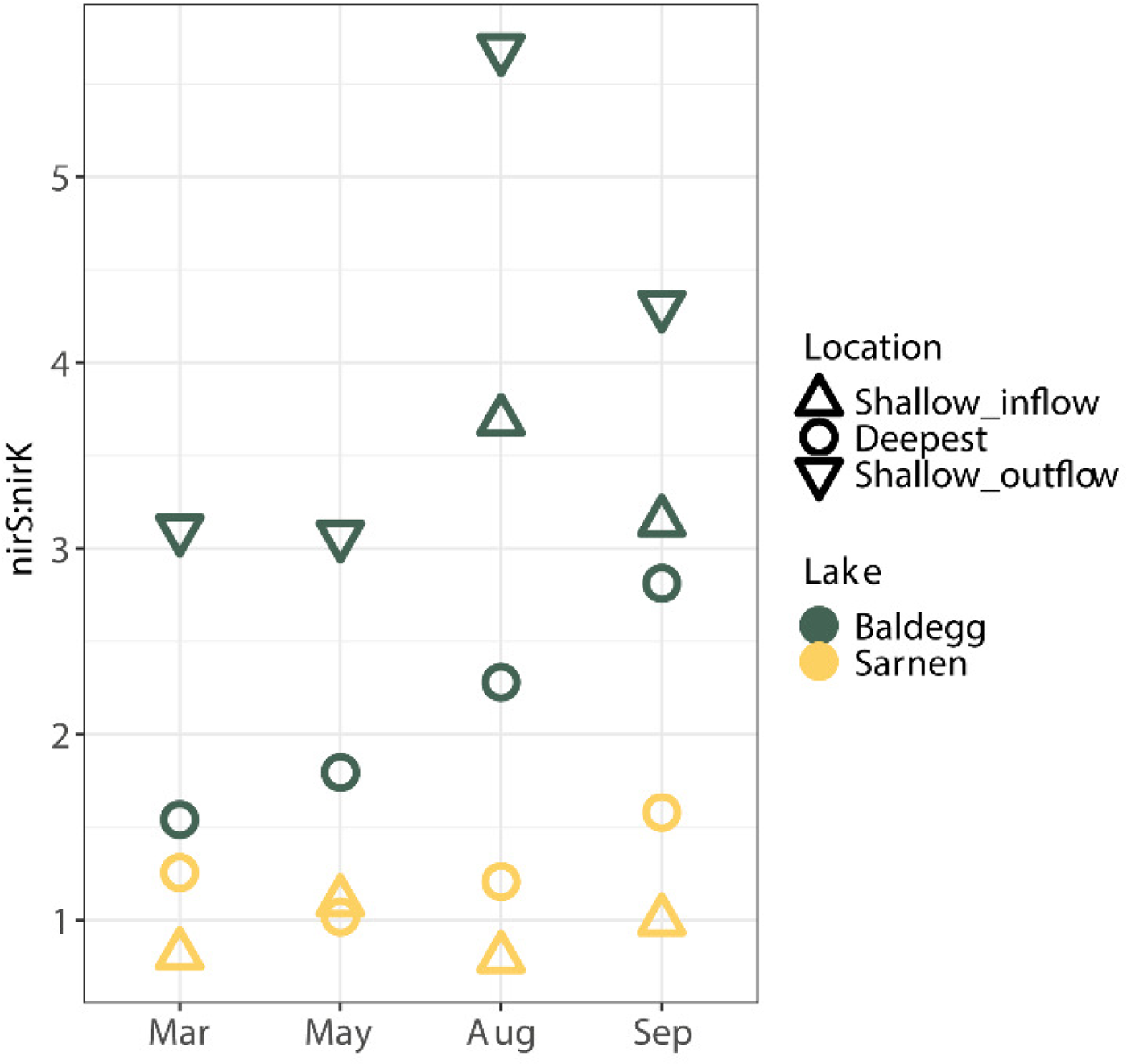
nirS:nirK ratio calculated from gpm values per Lake (Baldegg= green, Sarnen=yellow), Location (Shallow inflow: triangle, Deepest: plus, Shallow outflow: circle) and Month.

About 47% of the significant N transformation genes could be taxonomically assigned to at least class level. The results suggest that different key players dominate N cycling in each lake (Tab. 2). The overall most represented phyla involved in the N cycle were the *Gammaproteobacteria* (2950 gpm), followed by *Deltaproteobacteria* (1462 gpm), *Bacteroidetes* (720 gpm), and *Nitrospirae* (638 gpm). For Nitrification, the most distinct key players were the nitrite-oxidizing bacteria (NOB) *Nitrospirae* (300 gpm) in Lake Sarnen, which were present only in very low abundance in Lake Baldegg (9 gpm) (Tab. 2). Not only *nxrB* but also *amoABC* genes were assigned to *Nitrospirae* in Lake Sarnen. In contrast, the ammonia-oxidizing bacteria (AOB) *Nitrosomonadales* (*Gammaproteobacteria*) were represented in both Lakes. For the nitrate and nitrite reductase processes from ANRA, DNRA, and Dentrification_DNRA (NO_3_^-^ reduction), a phylogenetically diverse group of organisms appears to be involved (Tab. 2). In Lake Sarnen, the most abundant phyla carrying *nirK* were *Nitrospirae* (157 gpm) and *Chloroflexi* (113 gpm), whereas, in Lake Baldegg, this function was assigned mainly to *Gammaproteobacteria* (86 gpm) and *Bacteroidetes* (52 gpm) (Tab. 2). The most frequently assigned *nirS* phylum was *Gammaproteobacteria*, representing 448 gpm in Lake Baldegg and 211 gpm in Lake Sarnen, respectively. The most abundant phyla containing the nitrous oxide (*nosZ*) reducing denitrification genes in Lake Baldegg and Sarnen were *Bacteroidetes* (140 and 54 gpm) and *Gammaproteobacteria* (158 and 49 gpm). Nitrogenase genes in Lake Baldegg were often assigned to *Euryarchaeota* (173 gpm).

## Discussion

### Distinct microbial communities and porewater chemistry

The porewater chemistry in the studied eutrophic and oligotrophic lake sediments was correlated with clearly distinct microbial community composition. Interestingly, we observed that while the chemical sediment porewater signature was to a considerable extent seasonally driven ^20^, the microbial community composition was mainly affected by locational differences but relatively stable over time (Fig. 4, Fig. S2). The sediment porewater profiles of eutrophic Lake Baldegg indicated a high availability of N compounds (Fig. 2). Due to bottom water aeration, the O_2_ concentration remained high in this lake. The rapid depletion of O_2_, NO_3_^-^, NO_2_^-^ and SO_4_^2-^ and the increasing concentration of NH_4_^+^ with sediment depth indicated a high level of respiration activity, which is in line with previous reports ^20, 42, 62^. Lake Sarnen, in contrast, showed similar sediment porewater profiles as other oligotrophic lakes, e.g., Lakes Lucerne ^42^, Brienz, or Thun ^62^. However, compared to these lakes, the N compound concentrations in Lake Sarnen were lower, the O_2_ and SO_4_^2^ concentrations, and O_2_ penetration depth higher, which could be a sign of lower respiratory activity (Fig. 2). Overall, the porewater data confirmed that the lake sediments chosen for this study were suitable endmembers of the oligotrophic to the eutrophic spectrum of lakes with aerobic water columns in Switzerland.

Although the lakes share many microbial taxa, the alpha and beta diversity analysis nonetheless indicated a very distinct community composition for Lakes Baldegg and Sarnen (Fig. 4, S1 and S2). The most abundant phyla occurring in both lakes were *Proteobacteria*, *Bacteroidetes*, *Nitrospirae*, *Verrucomicrobia*, *Acidobacteria*, and *Chloroflexi* (Fig. 3), which is similar to microbial community compositions reported from other lake sediments ^25, 63–68^. Lake Sarnen showed a microbial community composition similar to other oligotrophic lakes, such as Lake Lucerne ^63^ or alpine lakes from the Henguan Mountains ^65^, in which a comparatively high *Nitrospirae* and *Alphaproteobacteria* abundance was reported. This similarity indicates that our results for the detailed analysis of functional potential may be to some extent generalizable. The significantly higher species richness in Lake Sarnen compared to Lake Baldegg is explained by Lake Sarnen harboring a greater number of fastidious oligotrophic microbes due to the low nutrient content. In contrast, the high nutrient abundance in Lake Baldegg favors a smaller number of competitive, copiotrophic microbes.

The microbial community composition changes with sediment depth are directly linked to the sediment’s redox cascade (Fig. S2A and B). Several specific phyla or classes show corresponding abundance patterns with sediment depth. For example, we found that *Nitrospirae* were most abundant in the surface sediment, especially in Lake Sarnen (Fig S3B). *Nitrospirae* is a known aerobic NOB occurring at low NH_4_^+^ concentrations ^69^, which goes herein with the low NH_4_^+^ and high O_2_ concentrations measured in Lake Sarnen. The same habitat preferences were reported for *Nitrososphaeria,* ammonia-oxidizing archaea (AOA) ^70^, which were only found in Lake Sarnen sediments. The absence of NOB and AOA in Lake Baldegg sediments can be explained by the fast O_2_ depletion and high NH_4_^+^ concentrations. In contrast, the *Deltaproteobacteria*, which includes many anaerobic sulfate-reducing bacteria ^71^, and *Methanomicrobia* (anaerobic Euryarchaeota) became more dominant with increasing sediment depth in Lake Baldegg (Fig. S3A). Similar trends in Lake Baldegg were reported in 2016 by ^63^. Seasonal changes in microbial community composition were only visible at the sediment surface, where the abundance of *Cyanobacteria* increased in August and September, mainly in Lake Baldegg (Fig. S3A) and less strongly in Lake Sarnen (Fig. S3B). This finding likely reflects the seasonal sedimentation of biomass from cyanobacterial blooms in the surface water in late spring and summer, which was also observed in the sediment traps ^20^.

We found the most distinct microbial community composition at the shallow inflow location of Lake Baldegg, which could be explained by high nutrient and organic matter input from the tributary. Another explanation for locational differences can be sediment focusing. Both lakes have rather steep shorelines (Fig. 1), which can lead to sediment movements towards the deeper parts of the lake in a process called sediment focusing ^72^. The overall lower locational differences in Lake Sarnen could be explained by the influence of its alpine river tributaries, which can have very high flow rates and transport large amounts of terrestrial material ^73^. During storm events, we measured an increase of sedimentation rates from ∼8 g m^-2^ d^-1^ to 556 g m^-2^ d^-1^ in June 2017 and 112 g m^-2^ d^-1^ in January 2018, at the deepest station ∼2 km from the inflow ^20^. We, therefore, suggest that the larger input of terrestrial OM and river sediments and its wide distribution in the lake contribute to lower locational niche diversity in Lake Sarnen compared to Lake Baldegg.

### Microbial nitrogen transformation potential

Our metagenomic analysis revealed that both lakes harbor microbial communities with the potential to perform almost all known N transformation processes (Fig. 8). The N gene composition and taxonomical data indicated locational differences in the microbial community composition and N transformation potential between and within both lakes. This is in agreement with previous observations of a local difference in denitrification rates in lake sediments ^25^. This finding emphasizes the importance of including locational differences of microbial community composition and function in lake sediment studies. In contrast to the system and location effects, we saw little evidence for seasonal variation in the sediment microbial communities or their N cycle potential. This lack of seasonal variation indicates that community assembly, together with its N transformation potential, plays out over longer time scales and does not adapt rapidly to minor seasonal environmental changes. This does not exclude short-term responses of activity, which can only be clarified by N transformation rate measurements, and applying methods that characterize the active microbial community, e.g. metatranscriptomics or -proteomics.

**FIGURE 8:**
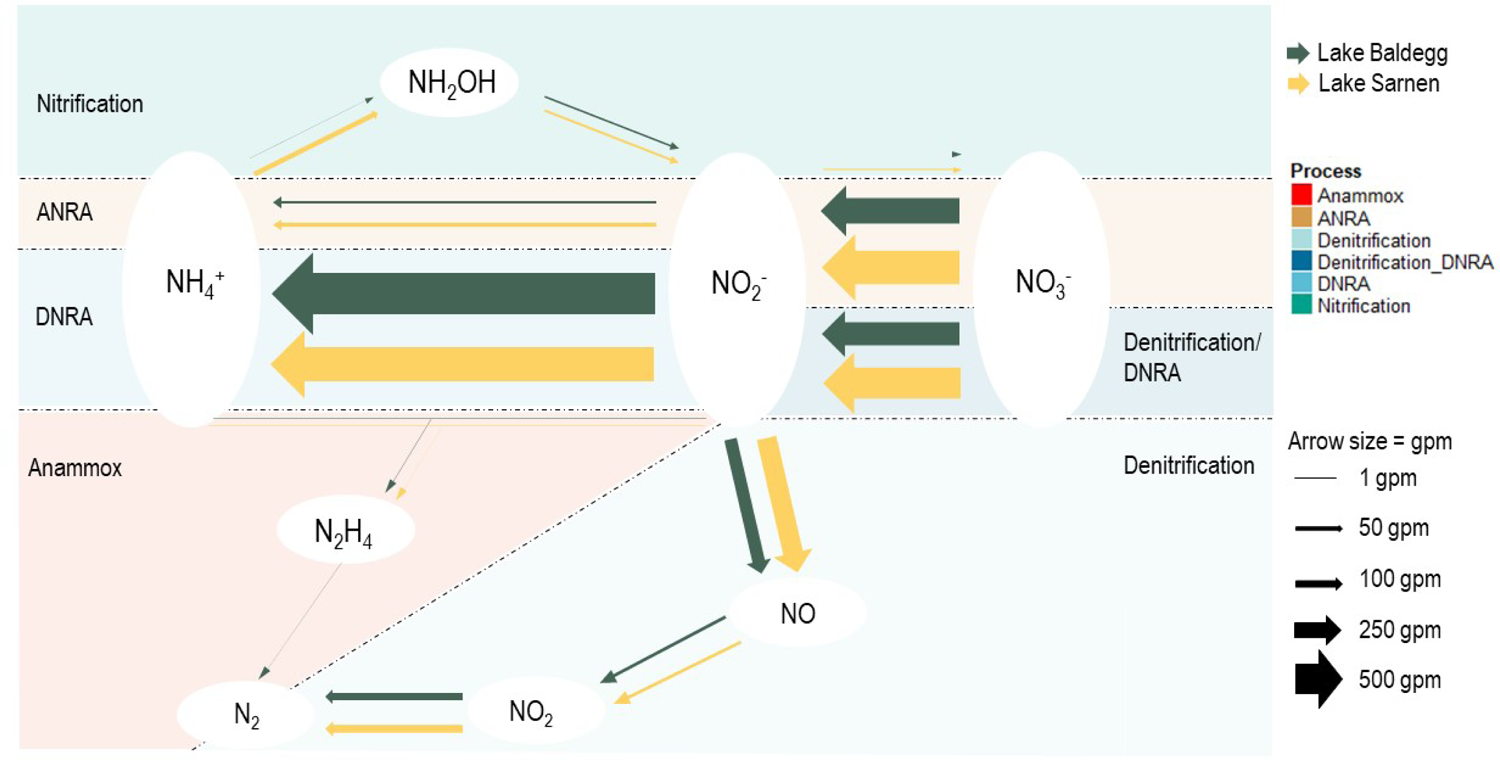
Conceptual model of the nitrogen transformation processes in Lake Baldegg (green arrows) and Lake Sarnen (yellow arrows). The arrow thickness indicates the average gpm per Lakes for the different transformation processes. The gene families are clustered in the different transformation processes: Nitrification (turquoise), ANRA (brown), Denitrification_DNRA nitrate reductase genes (dark blue), DNRA (bright blue), Denitrification (bright turquoise), Anammox (red) and Nitrogen (N_2_) fixation (N_fixation in black).

### Nitrogen inputs - organic N transformation and N2 fixation

The abundance of genes related to organic N transformation (N_degradation) was very high in both lakes, indicating that microbes can degrade organic N compounds in the OM input to produce N_r_ (Fig. 5). Another way for the ecosystem to get N_r_ is N_2_ fixation, and nitrogenase genes were found in all samples (Fig. 5). In the water column of lakes, N_2_ fixation is a well-known process ^3,22, 39, 74^. Despite our observation of seasonal cyanobacterial deposition in the microbial community data, cyanobacterial nitrogenase genes were not identified in the sediment of Lake Baldegg. The high N_2_ fixation gene abundance in Lake Baldegg was interestingly assigned mainly to methanogenic *Euryarchaeota* (*Methanomicrobiales*) and *Proteobacteria* (Tab. 2). Methanogenic *Euryarchaeota* were reported from wetland soils where they actively transcribed *nifH* genes ^75^. This leads to the assumption that Lake Baldegg’s sediment microbial community could respond to N scarcity. However, due to the high NH_4_^+^ concentration in the sediments, this is unlikely to represent an important N input at the moment, but more research would be required to verify this assumption.

### Nitrification

Lake Sarnen’s higher nitrification potential agrees with the higher O_2_ concentration and penetration depth, as it is an aerobic process. Similar results have been reported for river freshwater sediments previously ^76^. In Lake Baldegg, the shallow O_2_ penetration depth in the sediments likely constrains the development of an abundant nitrification community, which is in line with our rate measurements ^77^. Instead, nitrification dominates in the overlaying water in Lake Baldegg where O_2_ is more readily available ^77^. Lake Baldegg showed the highest abundance for *Nitrosomonadales* among identified AOB. This taxon is known for partial nitrification by encoding the oxidation step of NH_4_^+^ to NO_2_^-^ (*nxrB*), and can thrive under low O_2_ concentrations ^1,23^. *Nitrospirales,* well-known NOBs *^1,^*^39, 70, 76^, were the main taxa assigned to *nxrB* in Lake Sarnen (Table 2). Interestingly, *amoABC_B* genes were also assigned to *Nitrospirales* in Lake Sarnen. Recent studies reported *Nitrospira* species carrying all nitrification genes (*amo*, *hao,* and *nxr* genes) and being capable of complete nitrification, known as comammox ^30, 31, 78^. Comammox-*Nitrospira* were also reported from wetlands, agricultural and forest soil, wastewater treatment plants, and drinking water systems ^79^ and in lake sediments from Lake Superior ^25^. To test the hypothesis that Comammox*-Nitrospira* are abundant in Lake Sarnen, we mapped the bacterial *amoABC* genes against four comammox *Nitrospira* and two non-comammox *Nitrospira* genomes ^30, 78, 80, 81^. We found *amoABC* and *nxrB* genes from Lake Sarnen mapping to the Comammox-*Nitrospira* genomes but not those from Lake Baldegg, which supports our the hypothesis (Tables S5). Due to insufficient sequencing depth, we were unable to retrieve enough high-quality metagenome-assembled genomes (MAGs) to confirm the presence of comammox *Nitrospira*. Hence, at this time, we can not provide further confirmation whether comammox in Lake Sarnen sediments contributes to NH_4_^+^ oxidation.

**TABLE 2:**
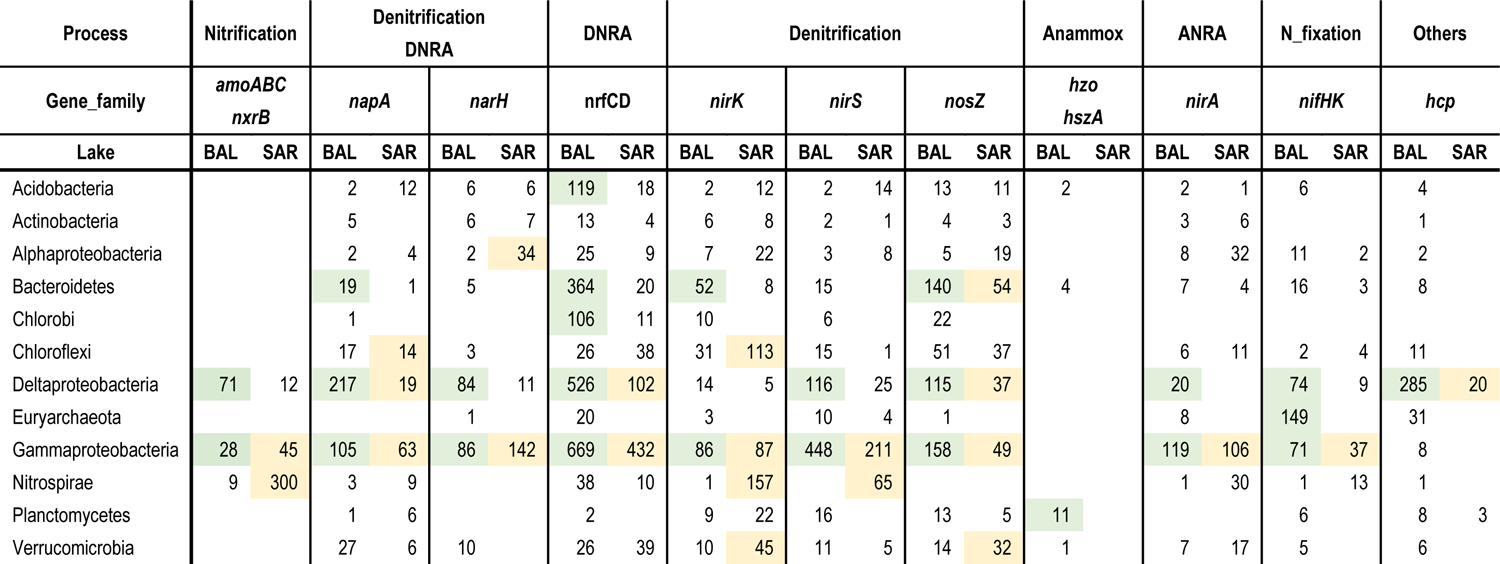
Overview of the mapping rates of the N removal genes to Kaiju database, to reveal possible key players (Top 12 Phyla) for the significantly different gene families, plus nirS. The numbers are the total sum of gpm per genes per Lake and Phyla. The most abundant Phyla are highlighted in green for Lake Baldegg and yellow Lake Sarnen.

### NO_3_^-^ and NO_2_^-^ reduction processes

Interestingly, we measured lower transformation rates for DNRA than for denitrification in both lakes ^77^, even though the gene abundance for DNRA was higher comparing to denitrification (Table S4). Many environmental drivers are playing a role in the competition between denitrification and DNRA. For example, higher input of labile organic carbon (C), increasing C:N ratio, and the ratio of electron donors to electron acceptors, and sulfide concentrations are all known to affect the balance between these processes ^18, 27, 39, 82–84^. It seems that many microbes in Lake Baldegg and Sarnen have the capacity to perform DNRA if the situation is favorable for this process, but generally rely more on denitrification under current conditions, probably because this process has a higher energy yield than DNRA ^39^. The more balanced ratio of denitrification to DNRA activity in Lake Sarnen measured by Callbeck et al.^77^ may be a consequence of the higher C:N ratios or the presumably higher sulfide levels from sulfate reduction, both of which can promote DNRA ^18, 39^. On the other hand, inhibition of DNRA due to high NO_3_^-^ concentrations has been observed ^82^, which could explain the low DNRA rates in Lake Baldegg. However, it is not yet clearly understood what conditions favor a switch to DNRA in lake sediments ^39^.

There are other processes potentially competing for NO_2_^-^, such as anammox. Callbeck et al. ^77^ measured low anammox rates compared to denitrification rates in the deepest station of Lake Baldegg, whereas in Lake Sarnen, the anammox rates in the surface sediment were as high as the denitrification rates. This is interesting, as we could find some but not the complete set of anammox genes in Lake Sarnen. This suggests a low abundance of anammox bacteria, which is confirmed by the low abundance of *Planctomycetes* (the phylum that contains all known anammox bacteria) for both lakes (Baldegg 0.01%, Lake Sarenen 0.1% from all 16S rRNA sequences). Low anammox bacterial abundance does not mean they are not active. This was reported from other aquatic ecosystems with high anammox rates and low *Planctomycetes* abundance ^11, 15, 85, 86^. In Lake Superior, sediments have deep O_2_ and NO_3_^-^ penetration depth, and anammox can contribute up to 50% N loss ^28^. Crowe et al. ^28^ stated that they predict that anammox might be an important N loss process for low productive freshwater environments such as high-latitude lakes. Higher sequencing depth of metagenomes and metatranscriptome would help to resolve the probably higher anammox potential and activity in the oligotrophic Lake Sarnen.

### Denitrification

The higher abundance of *nosZ* in Lake Baldegg, especially at the deep station (Fig. 6), is in good agreement with the higher denitrification rates measured by Callbeck et al. ^77^ and higher N removal rates reported by Müller et al. ^20^. Graf et al. ^29^ found that *nosZ* genes have a higher co-occurrence with *nirS* than *nirK*, thus *nirS* denitrifiers are more likely to do complete denitrification, as they also often co-occur with *nor* and *nosZ* genes. Our observation of a higher *nirS:nirK* ratio in Lake Baldegg compared to Lake Sarnen suggests that the capability for complete denitrification is more important in this lake. *nirK* denitrifiers are reported to respond differently to environmental gradients than *nirS* denitrifiers and occupy a different ecological niche ^29^. These results suggest that lakes with higher trophic status harbor denitrifiers with the *nirS gene and* the capacity for complete denitrification, while oligotrophic lakes provide an ecological niche for *nirK* denitrifiers. Thus the *nirS:nirK* ratio differs in lakes with different trophic states (Fig. 7).

### Nitrogen removal in a eutrophic and oligotrophic lake from the microbe’s perspective

Our data showed that even though our eutrophic and oligotrophic lakes have similar overall microbial N transformation potential, the evidence for a greater importance of nitrification (including comammox), DNRA, and anammox in Lake Sarnen collectively indicate a greater importance of internal N cycling via NH_4_^+^ in the oligotrophic Lake Sarnen. In addition, the key players involved in N processing differ significantly. Our findings are reflected in different N transformation rates and N removal efficiency in the studied lakes. Müller et al. ^20^ calculated a much higher N removal rate (NRR) (20+-6.6 g N m^-2^ yr^-1^) and N removal efficiency (NRE; 66%) for Lake Baldegg compared to Lake Sarnen with a NRR of 3.2+-4.2 g N m^-2^ yr^-1^ and NRE of 33%. In agreement, Callbeck et al. ^77^ measured more than 15 times higher denitrification rates in Lake Baldegg sediments (7.7 g N m^-2^ yr^-1^) than in Lake Sarnen (0.5 g N m^-2^ yr^-1^) in parallel to our study. Low denitrification activity in oligotrophic lakes is a known phenomenon, reported from the Laurentian Great Lakes ^25^. They stated that oligotrophic lakes might be on their microbial N activity threshold and can not denitrify more even with excess NO_3_^-^ concentrations. The higher denitrification rates in Lake Baldegg could be linked to the higher ratio of *nirS:nirK* denitrifiers in Lake Baldegg, which may harbor the potential for complete denitrification. The higher TOC input can also accelerate the microbial activity in the N cycle, as it acts as an electron donor for various N reduction pathways such as heterotrophic and chemolithoautotrophic denitrification ^1,18, 39^. Furthermore, we measured a higher NO_3_^-^ concentration in Lake Baldegg that enhanced denitrification while inhibiting DNRA ^87^. Both reasons could explain the lower denitrification activity in Lake Sarnen. The considerable diversity and abundance of N transformation genes in the sediment surface layer show the tight connection of the various processes as reported in other studies on the coupling of nitrification and denitrification in sediments ^19^. The combination of high TOC deposition, high NO_3_^-^ concentration, and artificial aeration of Lake Baldegg appears to create a good ecological niche for denitrifying bacteria. Lake Sarnen, in contrast, is less efficient in terms of N removal, but as an oligotrophic lake, nevertheless exports little N, while harboring a high microbial and macrobiological diversity with important ecosystem functions. We suggest that the survey of the microbial ecology can ostensibly inform our understanding of the lake biogeochemistry and vice versa.

## Conclusion

The microbial potential for N transformation in lakes appears to be strongly affected by trophic status as well as by the OM sediment deposition and N load from tributaries. Metagenomic data from the oligotrophic and eutrophic lake showed that microbial communities contained all N transformation genes, except the genes for the last step of the anammox process, which was not detected in Lake Sarnen. Furthermore, the overall N removal and the specific N transformation rates could be connected with the microbial N functional and taxonomical footprint. N transformation potential in lake sediments is clearly adapted to the environmental conditions (e.g., porewater gradient, OM input, etc.), and process responses to changing environmental conditions may thus, at least in the short term, depend on the properties of the established microbial communities. We suggest the *nirS:nirK* ratio of denitrifiers might indicate N removal efficiency and that oligotrophic lake sediments harbor higher microbial diversity.

## Acknowledgments

We want to thank the technical staff of Eawag, namely Patrick Kathriner, Karin Beck, Alois Zwyssig, Serge Robert and Nina Studhalter, for their help during fieldwork and in the analytical, biogeochemical and microbial laboratory. Thanks to Xinguo Han for his support in troubleshooting DNA/RNA extraction from the two lakes. We are grateful to the cantonal authorities Obwalden (Lake Sarnen) and ProNatura and Canton Lucerne (Lake Baldegg) for supporting our study by granting access to the lakes for sampling and permission to deploy sediment traps. We want to thank the genomic diversity center (GDC) and Euler Cluster at ETH for the bioinformatic support and access to their computational resources for the metagenomic analysis. We also want to thank Martin Schmid and Bernhard Wehrli for the fruitful discussions of the data. This work was financially supported by the SNF grant Regulation of Nitrogen turnover in Lakes (205321_169142).

## References

1. Kuypers, M. M. M.; Marchant, H. K.; Kartal, B. The Microbial Nitrogen-Cycling Network. Nat. Rev. Microbiol. 2018, 16 (5), 263–276. https://doi.org/10.1038/nrmicro.2018.9.

2. Isobe, K.; Ohte, N. Ecological Perspectives on Microbes Involved in N-Cycling. Microbes Environ. 2014, 29 (1), 4–16. https://doi.org/10.1264/jsme2.me13159.

3. Zehr, J. P.; Kudela, R. M. Nitrogen Cycle of the Open Ocean: From Genes to Ecosystems. Ann. Rev. Mar. Sci. 2011, 3, 197–225. https://doi.org/10.1146/annurev-marine-120709-142819.

4. Gruber, N.; Galloway, J. N. An Earth-System Perspective of the Global Nitrogen Cycle. Nature 2008, 451 (7176), 293–296. https://doi.org/10.1038/nature06592.

5. Seitzinger, S.; Harrison, J. A.; Böhlke, J. K.; Bouwman, A. F.; Lowrance, R.; Peterson, B.; Tobias, C.; Van Drecht, G. Denitrification across Landscapes and Waterscapes: A Synthesis. Ecol. Appl. 2006, 16 (6), 2064–2090. https://doi.org/10.1890/1051-0761(2006)016[2064:dalawa]2.0.co;2.

6. BAFU. Stickstoffflüsse in der Schweiz; UW-1018-D; BAFU, 2010.

7. Holland, E. A.; Braswell, B. H.; Sulzman, J.; Lamarque, J.-F. Nitrogen Deposition onto the United States and Western Europe: Synthesis of Observations and Models. Ecol. Appl. 2005, 15 (1), 38–57. https://doi.org/10.1890/03-5162.

8. Eawag: Swiss Federal Institute of Aquatic Science and Technology; Federal Office for the Environment (FOEN). NADUF – National long-term surveillance of Swiss rivers https://opendata.eawag.ch/group/naduf-national-long-term-surveillance-of-swiss-rivers (accessed Apr 10, 2018). https://doi.org/10.25678/00036W.

9. Billen, G.; Silvestre, M.; Grizzetti, B.; Leip, A.; Garnier, J.; Voss, M.; Howarth, R.; Bouraoui, F.; Lepistö, A.; Kortelainen, P.; Johnes, P.; Curtis, C.; Humborg, C.; Smedberg, E.; Kaste, Ø.; Ganeshram, R.; Beusen, A.; Lancelot, C. Nitrogen Flows from European Regional Watersheds to Coastal Marine Waters. The European Nitrogen Assessment. 2011, pp 271–297. https://doi.org/10.1017/cbo9780511976988.016.

10. Bernhardt, E. S. Ecology. Cleaner Lakes Are Dirtier Lakes. Science 2013, 342 (6155), 205–206. https://doi.org/10.1126/science.1245279.

11. Schubert, C. J.; Durisch-Kaiser, E.; Wehrli, B.; Thamdrup, B.; Lam, P.; Kuypers, M. M. M. Anaerobic Ammonium Oxidation in a Tropical Freshwater System (Lake Tanganyika). Environ. Microbiol. 2006, 8 (10), 1857–1863. https://doi.org/10.1111/j.1462-2920.2006.01074.x.

12. Finlay, J. C.; Small, G. E.; Sterner, R. W. Human Influences on Nitrogen Removal in Lakes. Science 2013, 342 (6155), 247–250. https://doi.org/10.1126/science.1242575.

13. Thamdrup, B. New Pathways and Processes in the Global Nitrogen Cycle. Annual Review of Ecology Evolution and Systematics 2012, 43, 407–428. https://doi.org/10.1146/annurev-ecolsys-102710-145048.

14. Penton, C. R.; Devol, A. H.; Tiedje, J. M. Molecular Evidence for the Broad Distribution of Anaerobic Ammonium-Oxidizing Bacteria in Freshwater and Marine Sediments. Appl. Environ. Microbiol. 2006, 72 (10), 6829–6832. https://doi.org/10.1128/AEM.01254-06.

15. Kuypers, M. M. M.; Sliekers, A. O.; Lavik, G.; Schmid, M.; Jørgensen, B. B.; Kuenen, J. G.; Sinninghe Damsté, J. S.; Strous, M.; Jetten, M. S. M. Anaerobic Ammonium Oxidation by Anammox Bacteria in the Black Sea. Nature 2003, 422 (6932), 608–611. https://doi.org/10.1038/nature01472.

16. Dalsgaard, T.; Canfield, D. E.; Petersen, J.; Thamdrup, B.; Acuña-González, N2 Production by the Anammox Reaction in the Anoxic Water Column of Golfo Dulce, Costa Rica. Nature 2003, 422 (6932), 606–608. https://doi.org/10.1038/nature01526.

17. Hirsch, M. D.; Long, Z. T.; Song, B. Anammox Bacterial Diversity in Various Aquatic Ecosystems Based on the Detection of Hydrazine Oxidase Genes (hzoA/hzoB). Microb. Ecol. 2011, 61 (2), 264–276. https://doi.org/10.1007/s00248-010-9743-1.

18. Kraft, B.; Tegetmeyer, H. E.; Sharma, R.; Klotz, M. G.; Ferdelman, T. G.; Hettich, R. L.; Geelhoed, J. S.; Strous, M. Nitrogen Cycling. The Environmental Controls That Govern the End Product of Bacterial Nitrate Respiration. Science 2014, 345 (6197), 676–679. https://doi.org/10.1126/science.1254070.

19. Marchant, H. K.; Holtappels, M.; Lavik, G.; Ahmerkamp, S.; Winter, C.; Kuypers, M. M. M. Coupled Nitrification–denitrification Leads to Extensive N Loss in Subtidal Permeable Sediments. Limnol. Oceanogr. 2016, 61 (3), 1033–1048. https://doi.org/10.1002/lno.10271.

20. Müller, B.; Thoma, R.; Baumann, K. B. L.; Callbeck, C. M.; Schubert, C. J. Nitrogen Removal Processes in Lakes of Different Trophic States from on-Site Measurements and Historic Data. Aquat. Sci. 2021, 83 (2). https://doi.org/10.1007/s00027-021-00795-7.

21. Kalvelage, T.; Lavik, G.; Lam, P.; Contreras, S.; Arteaga, L.; Löscher, C. R.; Oschlies, A.; Paulmier, A.; Stramma, L.; Kuypers, M. M. M. Nitrogen Cycling Driven by Organic Matter Export in the South Pacific Oxygen Minimum Zone. Nature Geoscience. 2013, pp 228–234. https://doi.org/10.1038/ngeo1739.

22. Callbeck, C. M.; Ehrenfels, B.; Baumann, K. B. L.; Wehrli, B.; Schubert, C. J. Anoxic Chlorophyll Maximum Enhances Local Organic Matter Remineralization and Nitrogen Loss in Lake Tanganyika. Nat. Commun. 2021, 12 (1), 830. https://doi.org/10.1038/s41467-021-21115-5.

23. Bristow, L. A.; Dalsgaard, T.; Tiano, L.; Mills, D. B.; Bertagnolli, A. D.; Wright, J.; Hallam, S. J.; Ulloa, O.; Canfield, D. E.; Revsbech, N. P.; Thamdrup, B. Ammonium and Nitrite Oxidation at Nanomolar Oxygen Concentrations in Oxygen Minimum Zone Waters. Proc. Natl. Acad. Sci. U. S. A. 2016, 113 (38), 10601–10606. https://doi.org/10.1073/pnas.1600359113.

24. Rissanen, A. J.; Tiirola, M.; Hietanen, S.; Ojala, A. Interlake Variation and Environmental Controls of Denitrification across Different Geographical Scales. Aquat. Microb. Ecol. 2013, 69 (1), 1–16. https://doi.org/10.3354/ame01619.

25. Small, G. E.; Finlay, J. C.; McKay, R. M. L.; Rozmarynowycz, M. J.; Brovold, S.; Bullerjahn, G. S.; Spokas, K.; Sterner, R. W. Large Differences in Potential Denitrification and Sediment Microbial Communities across the Laurentian Great Lakes. Biogeochemistry 2016, 128 (3), 353–368. https://doi.org/10.1007/s10533-016-0212-x.

26. Tu, Q.; He, Z.; Wu, L.; Xue, K.; Xie, G.; Chain, P.; Reich, P. B.; Hobbie, S. E.; Zhou, J. Metagenomic Reconstruction of Nitrogen Cycling Pathways in a CO2-Enriched Grassland Ecosystem. Soil Biol. Biochem. 2017, 106, 99–108. https://doi.org/10.1016/j.soilbio.2016.12.017.

27. Giblin, A.; Tobias, C.; Song, B.; Weston, N.; Banta, G.; Rivera-Monroy, V. The Importance of Dissimilatory Nitrate Reduction to Ammonium (DNRA) in the Nitrogen Cycle of Coastal Ecosystems. Oceanography. 2013, pp 124–131. https://doi.org/10.5670/oceanog.2013.54.

28. Crowe, S. A.; Treusch, A. H.; Forth, M.; Li, J.; Magen, C.; Canfield, D. E.; Thamdrup, B.; Katsev, S. Novel Anammox Bacteria and Nitrogen Loss from Lake Superior. Scientific Reports. 2017. https://doi.org/10.1038/s41598-017-12270-1.

29. Graf, D. R. H.; Jones, C. M.; Hallin, S. Intergenomic Comparisons Highlight Modularity of the Denitrification Pathway and Underpin the Importance of Community Structure for N2O Emissions. PLoS ONE. 2014, p e114118. https://doi.org/10.1371/journal.pone.0114118.

30. Daims, H.; Lebedeva, E. V.; Pjevac, P.; Han, P.; Herbold, C.; Albertsen, M.; Jehmlich, N.; Palatinszky, M.; Vierheilig, J.; Bulaev, A.; Kirkegaard, R. H.; von Bergen, M.; Rattei, T.; Bendinger, B.; Nielsen, P. H.; Wagner, M. Complete Nitrification by Nitrospira Bacteria. Nature 2015, 528 (7583), 504–509. https://doi.org/10.1038/nature16461.

31. Koch, H.; van Kessel, M. A. H. J.; Lücker, S. Complete Nitrification: Insights into the Ecophysiology of Comammox Nitrospira. Appl. Microbiol. Biotechnol. 2019, 103 (1), 177–189. https://doi.org/10.1007/s00253-018-9486-3.

32. Tu, Q.; Lin, L.; Cheng, L.; Deng, Y.; He, Z. NCycDB: A Curated Integrative Database for Fast and Accurate Metagenomic Profiling of Nitrogen Cycling Genes. Bioinformatics 2019, 35 (6), 1040–1048. https://doi.org/10.1093/bioinformatics/bty741.

33. Fierer, N.; Lauber, C. L.; Ramirez, K. S.; Zaneveld, J.; Bradford, M. A.; Knight, R. Comparative Metagenomic, Phylogenetic and Physiological Analyses of Soil Microbial Communities across Nitrogen Gradients. ISME J. 2012, 6 (5), 1007–1017. https://doi.org/10.1038/ismej.2011.159.

34. Nelson, M. B.; Martiny, A. C.; Martiny, J. B. H. Global Biogeography of Microbial Nitrogen-Cycling Traits in Soil. Proceedings of the National Academy of Sciences. 2016, pp 8033–8040. https://doi.org/10.1073/pnas.1601070113.

35. Orellana, L. H.; Rodriguez-R, L. M.; Higgins, S.; Chee-Sanford, J. C.; Sanford, R. A.; Ritalahti, K. M.; Löffler, F. E.; Konstantinidis, K. T. Detecting Nitrous Oxide Reductase (NosZ) Genes in Soil Metagenomes: Method Development and Implications for the Nitrogen Cycle. MBio 2014, 5 (3), e01193–14. https://doi.org/10.1128/mBio.01193-14.

36. Broadbent, A. A. D.; Snell, H. S. K.; Michas, A.; Pritchard, W. J.; Newbold, L.; Cordero, I.; Goodall, T.; Schallhart, N.; Kaufmann, R.; Griffiths, R. I.; Schloter, M.; Bahn, M.; Bardgett, R. D. Climate Change Alters Temporal Dynamics of Alpine Soil Microbial Functioning and Biogeochemical Cycling via Earlier Snowmelt. ISME J. 2021. https://doi.org/10.1038/s41396-021-00922-0.

37. Qi, D.; Feng, F.; Fu, Y.; Ji, X.; Liu, X. Effects of Soil Microbes on Forest Recovery to Climax Community through the Regulation of Nitrogen Cycling. Forests 2020, 11 (10), 1027. https://doi.org/10.3390/f11101027.

38. Lüke, C.; Speth, D. R.; Kox, M. A. R.; Villanueva, L.; Jetten, M. S. M. Metagenomic Analysis of Nitrogen and Methane Cycling in the Arabian Sea Oxygen Minimum Zone. PeerJ 2016, 4, e1924. https://doi.org/10.7717/peerj.1924.

39. Damashek, J.; Francis, C. A. Microbial Nitrogen Cycling in Estuaries: From Genes to Ecosystem Processes. Estuaries Coast. 2018, 41 (3), 626–660. https://doi.org/10.1007/s12237-017-0306-2.

40. Gächter, R.; Wehrli, B. Ten Years of Artificial Mixing and Oxygenation: No Effect on the Internal Phosphorus Loading of Two Eutrophic Lakes. Environ. Sci. Technol. 1998, 32 (23), 3659–3665. https://doi.org/10.1021/es980418l.

41. Steinsberger, T.; Müller, B.; Gerber, C.; Shafei, B.; Schmid, M. Modeling Sediment Oxygen Demand in a Highly Productive Lake under Various Trophic Scenarios. PLoS One 2019, 14 (10), e0222318. https://doi.org/10.1371/journal.pone.0222318.

42. Fiskal, A.; Deng, L.; Michel, A.; Eickenbusch, P.; Han, X.; Lagostina, L.; Zhu, R.; Sander, M.; Schroth, M. H.; Bernasconi, S. M.; Dubois, N.; Lever, M. A. Effects of Eutrophication on Sedimentary Organic Carbon Cycling in Five Temperate Lakes. Biogeosci. Discuss. 2019, 1–35. https://doi.org/10.5194/bg-2019-108.

43. Wickham, H. ggplot2: ggplot2. Wiley Interdiscip. Rev. Comput. Stat. 2011, 3 (2), 180–185. https://doi.org/10.1002/wics.147.

44. R Core Team. R: A language and environment for statistical computing https://www.r-project.org/ (accessed Jan 9, 2017).

45. Callahan, B. J.; McMurdie, P. J.; Rosen, M. J.; Han, A. W.; Johnson, A. J. A.; Holmes, S. P. DADA2: High-Resolution Sample Inference from Illumina Amplicon Data. Nat. Methods 2016, 13 (7), 581–583. https://doi.org/10.1038/nmeth.3869.

46. Callahan, B. DADA2 Pipeline Tutorial https://benjjneb.github.io/dada2/tutorial.html (accessed Jan 3, 2020).

47. Oksanen, J.; Kindt, R.; Legendre, P.; O’Hara, B.; Stevens, M. H. H.; Oksanen, M. J.; Suggests, M. The Vegan Package. Community ecology package 2007, 10 (631-637), 719.

48. McMurdie, P. J.; Holmes, S. Phyloseq: An R Package for Reproducible Interactive Analysis and Graphics of Microbiome Census Data. PLoS One 2013, 8 (4), e61217. https://doi.org/10.1371/journal.pone.0061217.

49. Ssekagiri, A.; Sloan, W. T.; Ijaz, U. Z. microbiomeSeq: An R Package for Microbial Community Analysis in an Environmental Context. 2018.

50. Andrews, S.; Others. FastQC: A Quality Control Tool for High Throughput Sequence Data. Babraham Bioinformatics, Babraham Institute, Cambridge, United Kingdom 2010.

51. Schmieder, R.; Edwards, R. Quality Control and Preprocessing of Metagenomic Datasets. Bioinformatics 2011, 27 (6), 863–864. https://doi.org/10.1093/bioinformatics/btr026.

52. Li, D.; Liu, C.-M.; Luo, R.; Sadakane, K.; Lam, T.-W. MEGAHIT: An Ultra-Fast Single-Node Solution for Large and Complex Metagenomics Assembly via Succinct de Bruijn Graph. Bioinformatics 2015, 31 (10), 1674–1676. https://doi.org/10.1093/bioinformatics/btv033.

53. Bushnell, B. BBMap: A Fast, Accurate, Splice-Aware Aligner; LBNL-7065E; Lawrence Berkeley National Lab. (LBNL), Berkeley, CA (United States), 2014.

54. Tarasov, A.; Vilella, A. J.; Cuppen, E.; Nijman, I. J.; Prins, P. Sambamba: Fast Processing of NGS Alignment Formats. Bioinformatics 2015, 31 (12), 2032–2034. https://doi.org/10.1093/bioinformatics/btv098.

55. Li, H.; Handsaker, B.; Wysoker, A.; Fennell, T.; Ruan, J.; Homer, N.; Marth, G.; Abecasis, G.; Durbin, R.; 1000 Genome Project Data Processing Subgroup. The Sequence Alignment/Map Format and SAMtools. Bioinformatics 2009, 25 (16), 2078–2079. https://doi.org/10.1093/bioinformatics/btp352.

56. Liao, Y.; Smyth, G. K.; Shi, W. featureCounts: An Efficient General Purpose Program for Assigning Sequence Reads to Genomic Features. Bioinformatics 2014, 30 (7), 923–930. https://doi.org/10.1093/bioinformatics/btt656.

57. Seemann, T. Prokka: Rapid Prokaryotic Genome Annotation. Bioinformatics 2014, 30 (14), 2068–2069. https://doi.org/10.1093/bioinformatics/btu153.

58. Love, M.; Anders, S.; Huber, W. Differential Analysis of Count Data—the DESeq2 Package. Genome Biol. 2014, 15 (550), 10–1186.

59. Love, M. I.; Anders, S.; Huber, W. Analyzing RNA-seq data with DESeq2 http://bioconductor.org/packages/release/bioc/vignettes/DESeq2/inst/doc/DESeq2.html (accessed Jan 6, 2019).

60. Gu, Z.; Eils, R.; Schlesner, M. Complex Heatmaps Reveal Patterns and Correlations in Multidimensional Genomic Data. Bioinformatics 2016, 32 (18), 2847–2849. https://doi.org/10.1093/bioinformatics/btw313.

61. Menzel, P.; Ng, K. L.; Krogh, A. Fast and Sensitive Taxonomic Classification for Metagenomics with Kaiju. Nat. Commun. 2016, 7, 11257. https://doi.org/10.1038/ncomms11257.

62. Steinsberger, T.; Schwefel, R.; Wüest, A.; Müller, B. Hypolimnetic Oxygen Depletion Rates in Deep Lakes: Effects of Trophic State and Organic Matter Accumulation. Limnol. Oceanogr. 2020, 65 (12), 3128–3138. https://doi.org/10.1002/lno.11578.

63. Han, X.; Schubert, C. J.; Fiskal, A.; Dubois, N.; Lever, M. A. Eutrophication as a Driver of Microbial Community Structure in Lake Sediments. Environ. Microbiol. 2020, 22 (8), 3446–3462. https://doi.org/10.1111/1462-2920.15115.

64. Wang, N. F.; Zhang, T.; Yang, X.; Wang, S.; Yu, Y.; Dong, L. L.; Guo, Y. D.; Ma, Y. X.; Zang, J. Y. Diversity and Composition of Bacterial Community in Soils and Lake Sediments from an Arctic Lake Area. Front. Microbiol. 2016, 7, 1170. https://doi.org/10.3389/fmicb.2016.01170.

65. Liao, B.; Yan, X.; Zhang, J.; Chen, M.; Li, Y.; Huang, J.; Lei, M.; He, H.; Wang, J. Microbial Community Composition in Alpine Lake Sediments from the Hengduan Mountains. Microbiologyopen 2019, 8 (9), e00832. https://doi.org/10.1002/mbo3.832.

66. Rissanen, A. J.; Peura, S.; Mpamah, P. A.; Taipale, S.; Tiirola, M.; Biasi, C.; Mäki, A.; Nykänen, H. Vertical Stratification of Bacteria and Archaea in Sediments of a Small Boreal Humic Lake. FEMS Microbiol. Lett. 2019, 366 (5). https://doi.org/10.1093/femsle/fnz044.

67. Huang, W.; Chen, X.; Jiang, X.; Zheng, B. Characterization of Sediment Bacterial Communities in Plain Lakes with Different Trophic Statuses. MicrobiologyOpen 2017, 6 (5). https://doi.org/10.1002/mbo3.503.

68. Zhang, L.; Zhao, T.; Wang, Q.; Li, L.; Shen, T.; Gao, G. Bacterial Community Composition in Aquatic and Sediment Samples with Spatiotemporal Dynamics in Large, Shallow, Eutrophic Lake Chaohu, China. J. Freshw. Ecol. 2019, 34 (1), 575–589. https://doi.org/10.1080/02705060.2019.1635536.

69. Herber, J.; Klotz, F.; Frommeyer, B.; Weis, S.; Straile, D.; Kolar, A.; Sikorski, J.; Egert, M.; Dannenmann, M.; Pester, M. A Single Thaumarchaeon Drives Nitrification in Deep Oligotrophic Lake Constance. Environ. Microbiol. 2020, 22 (1), 212–228. https://doi.org/10.1111/1462-2920.14840.

70. Pester, M.; Schleper, C.; Wagner, M. The Thaumarchaeota: An Emerging View of Their Phylogeny and Ecophysiology. Curr. Opin. Microbiol. 2011, 14 (3), 300–306. https://doi.org/10.1016/j.mib.2011.04.007.

71. Wörner, S.; Pester, M. The Active Sulfate-Reducing Microbial Community in Littoral Sediment of Oligotrophic Lake Constance. Front. Microbiol. 2019, 10, 247. https://doi.org/10.3389/fmicb.2019.00247.

72. Blais, J. M.; Kalff, J. The Influence of Lake Morphometry on Sediment Focusing. Limnol. Oceanogr. 1995, 40 (3), 582–588. https://doi.org/10.4319/lo.1995.40.3.0582.

73. Monecke, K.; Anselmetti, F. S.; Becker, A.; Sturm, M.; Giardini, D. The Record of Historic Earthquakes in Lake Sediments of Central Switzerland. Tectonophysics 2004, 394 (1), 21–40. https://doi.org/10.1016/j.tecto.2004.07.053.

74. Beversdorf, L. J.; Miller, T. R.; McMahon, K. D. The Role of Nitrogen Fixation in Cyanobacterial Bloom Toxicity in a Temperate, Eutrophic Lake. PLoS One 2013, 8 (2), e56103. https://doi.org/10.1371/journal.pone.0056103.

75. Bae, H.-S.; Morrison, E.; Chanton, J. P.; Ogram, A. Methanogens Are Major Contributors to Nitrogen Fixation in Soils of the Florida Everglades. Appl. Environ. Microbiol. 2018, 84 (7). https://doi.org/10.1128/AEM.02222-17.

76. Altmann, D.; Stief, P.; Amann, R.; De Beer, D.; Schramm, A. In Situ Distribution and Activity of Nitrifying Bacteria in Freshwater Sediment. Environ. Microbiol. 2003, 5 (9), 798–803.https://doi.org/10.1046/j.1469-2920.2003.00469.x.

77. Callbeck, C. M.; Baumann, K. B. L.; Thoma, R.; Müller, B.; Bürgmann, H.; Schubert, C. J. Freshwater Sediment Nitrogen Cycling Fueled by Boundary Layer Organic Nitrogen Remineralization to Ammonium. 2021.

78. van Kessel, M. A. H. J.; Speth, D. R.; Albertsen, M.; Nielsen, P. H.; Op den Camp, H. J. M.; Kartal, B.; Jetten, M. S. M.; Lücker, S. Complete Nitrification by a Single Microorganism. Nature 2015, 528 (7583), 555–559. https://doi.org/10.1038/nature16459.

79. Sun, D.; Tang, X.; Zhao, M.; Zhang, Z.; Hou, L.; Liu, M.; Wang, B.; Klümper, U.; Han, P. Distribution and Diversity of Comammox Nitrospira in Coastal Wetlands of China. Front. Microbiol. 2020, 11, 589268. https://doi.org/10.3389/fmicb.2020.589268.

80. Pinto, A. J.; Marcus, D. N.; Ijaz, U. Z.; Bautista-de Lose Santos, Q. M.; Dick, G. J.; Raskin, L. Metagenomic Evidence for the Presence of Comammox Nitrospira-Like Bacteria in a Drinking Water System. mSphere 2016, 1 (1). https://doi.org/10.1128/mSphere.00054-15.

81. Sakoula, D.; Nowka, B.; Spieck, E.; Daims, H.; Lücker, S. The Draft Genome Sequence of “Nitrospira Lenta” Strain BS10, a Nitrite Oxidizing Bacterium Isolated from Activated Sludge. Stand. Genomic Sci. 2018, 13 (1), 32. https://doi.org/10.1186/s40793-018-0338-7.

82. Heo, H.; Kwon, M.; Song, B.; Yoon, S. Involvement of NO3− in Ecophysiological Regulation of Dissimilatory Nitrate/Nitrite Reduction to Ammonium (DNRA) Is Implied by Physiological Characterization of Soil DNRA Bacteria Isolated via a Colorimetric Screening Method. Appl. Environ. Microbiol. 2020. https://doi.org/10.1128/AEM.01054-20.

83. Bu, C.; Wang, Y.; Ge, C.; Ahmad, H. A.; Gao, B.; Ni, S.-Q. Dissimilatory Nitrate Reduction to Ammonium in the Yellow River Estuary: Rates, Abundance, and Community Diversity. Sci. Rep. 2017, 7 (1), 6830. https://doi.org/10.1038/s41598-017-06404-8.

84. Burgin, A. J.; Hamilton, S. K. Have We Overemphasized the Role of Denitrification in Aquatic Ecosystems? A Review of Nitrate Removal Pathways. Front. Ecol. Environ. 2007, 5 (2), 89–96. https://doi.org/10.1890/1540-9295(2007)5[89:hwotro]2.0.co;2.

85. Pollet, T.; Humbert, J.-F.; Tadonléké, R. D. Planctomycetes in Lakes: Poor or Strong Competitors for Phosphorus? Appl. Environ. Microbiol. 2014, 80 (3), 819–828. https://doi.org/10.1128/AEM.02824-13.

86. Tadonléké, R. D. Strong Coupling between Natural Planctomycetes and Changes in the Quality of Dissolved Organic Matter in Freshwater Samples. FEMS Microbiol. Ecol. 2007, 59 (3), 543–555. https://doi.org/10.1111/j.1574-6941.2006.00222.x.

87. Hardison, A. K.; Algar, C. K.; Giblin, A. E.; Rich, J. J. Influence of Organic Carbon and Nitrate Loading on Partitioning between Dissimilatory Nitrate Reduction to Ammonium (DNRA) and N2 Production. Geochim. Cosmochim. Acta 2015, 164, 146–160. https://doi.org/10.1016/j.gca.2015.04.049.

